# Pancreatic cancer fibrosis activates protumorigenic Schwann cells through a nuclear mechanosensing mechanism

**DOI:** 10.64898/2026.04.21.719930

**Authors:** Pavel Stupakov, Golbahar Sadatrezaei, Ines Velazquez Quesada, Lillian A. Boe, Chun-Hao Chen, Francesca Gaino, Efsevia Vakiani, Ihsan Ekin Demir, Boris Reva, Bojana Gligorijevic, Richard J. Wong, Sylvie Deborde

## Abstract

**Background:** Fibrosis and tumor innervation are two features of the tumor microenvironment (TME) that contribute directly to the lethality of pancreatic ductal adenocarcinoma (PDAC), but their potential interactions have not been explored. Moreover, although it is known that activated Schwann cells (SCs) stimulate cancer cell invasion, it remains unclear how SCs are activated.

**Objective:** We determined how SCs are activated in the pancreatic fibrotic microenvironment.

**Design:** The correlation between physical features of the microenvironment and SC activation was assessed in human patient samples and in mice by SC c-Jun phosphorylation monitoring, atomic force microscopy and multiphoton live imaging. Several in vitro models in which forces were applied to SCs expressing a reporter for c-Jun phosphorylation and RNA-Seq analysis were used to decipher the cellular and molecular mechanisms of SC activation.

**Results:** Nerves surrounded by stiff stroma present higher SC activation. Intravital imaging shows a matrix dependent SC activation. Mechanical forces on SCs induce c-Jun phosphorylation in SCs in a non-canonical manner that involves a nuclear sensing machinery with the proinflammatory enzyme Phospholipase A2.

**Conclusion:** Fibrosis enhances the protumorigenic impact of innervation by activating SCs via a mechanism in which nuclear compression triggers non-canonical activation of the AP-1 transcription factor complex. Pancreatic fibrosis alone, without cancer cells, is sufficient to activate SCs, suggesting this mechanism may be common across non-malignant pancreatic diseases. Notably, SCs are more sensitive to mechanical activation than PDAC cells. These findings reveal TME interactions that may guide future microenvironment-targeted PDAC therapies.

**What is already known on this topic:** The pancreatic cancer tumor microenvironment is highly innervated and fibrotic, two components of the tumor microenvironment that regulate tumorigenesis. How they impact each other is unknown. Schwann cells have emerged as a significant protumorigenic player, but the triggers of Schwann cell activation remain undefined.

**What this study adds:** We establish that fibrosis induces Schwann cell activation and characterize the mechanism by which it occurs. We uncovered a mechanical mode of action that deforms nuclear membrane and activates c-Jun in Schwann cells, which contradicts the traditional view of c-Jun activation through a stimulus detected at the plasma membrane.

**How this study might affect research, practice or policy:** This study provides a better understanding of the biology of pancreatic ductal adenocarcinoma and supports the development of novel precision therapies that target the fibrotic microenvironment to impact the protumorigenic effect of tumor innervation.

## INTRODUCTION

Pancreatic ductal adenocarcinoma (PDAC) is one of the most lethal human solid malignancies, with a 5-year survival rate of 12.8% and rising incidence rates[1] Current treatments are inadequate despite a growing understanding of the biology of PDAC cells and components of the tumor microenvironment (TME) that promote tumor progression[2] Numerous studies have characterized the role of various pancreatic TME components on cancer cells, but how the pro-tumorigenic components modulate each other to affect the malignancy is unclear. Elucidating these relationships might provide new directions for disease treatment. Fibrosis and innervation are both features of the PDAC microenvironment, and have been reported to predict clinical outcomes.[3] Fibrosis is characterized by an accumulation of collagen and fibronectin which increases tumor stiffness and contributes to PDAC malignancy by forming a physical barrier that hinders immune cell infiltration and drug delivery. It can enhance the invasive properties of cancer cells by creating mechanical stress which induces phenotypic changes and affects migration.[4] However, the impact of fibrosis on other TME cell phenotypes remains unknown.

Pancreatic tumors are densely innervated, and nerves contribute to growth and metastasis of PDAC[5–7] and other cancer types.[8,9] Cancer cells can invade nerves, a process known as perineural invasion (PNI). PNI facilitates cancer cell migration from the primary tumor and can explain local recurrence after resection with seemingly clear margins. Schwann cells (SCs) are the most abundant cells in nerves and are actively involved in tumor spread, both in PDAC[10– 12] and in other cancer types.[13,14] SCs appear in PDAC before the onset of cancer invasion[15] and reprogram into nerve repair-like SCs.[10] Reprogrammed SCs interact dynamically with cancer cells by forming tracks that promote tumor invasion[11], support cancer-associated neuronal remodeling[16], induce malignant phenotypic switches in PDAC cancer cells and fibroblasts[17], and are associated with worse prognosis in PDAC.[11,17] SC reprogramming relies on the stress-related transcription factor c-Jun, canonically activated via phosphorylation through MAPK/JNK. However, the mechanism that triggers Schwann cell c-Jun activity in PDAC remains unknown.

Here, we study the interplay between fibrosis and the SCs of nerves. Monitoring of c-Jun phosphorylation in SCs in human specimens, as well as *in vitro* and live imaging assays in mice, reveals that phosphorylation is induced by mechanical forces through the activation of the nuclear sensor cytosolic Phospholipase A2 (cPLA2). We show that the stiffness caused by fibrosis can induce SC reprogramming by activating the transcription factor c-Jun in a non-canonical pathway. Our findings demonstrate that the fibrotic microenvironment of PDAC affects SC reprogramming and thus promotes PDAC malignancy through the diverse protumorigenic functions of SCs, including the process of perineural invasion.

## MATERIAL & METHODS

### Patient materials

Written informed consent was obtained from each patient. The studies were conducted in accordance with recognized ethical guidelines (e.g., Declaration of Helsinki, CIOMS, Belmont Report, U.S. Common Rule) and approved by the MSK Institutional Review Board (IRB #: 15-052)

### Mice

All experiments with mice were performed in accordance with an institutional animal care and use committee (IACUC)–approved protocol at Memorial Sloan Kettering Cancer Center (MSKCC). Experiments with intravital imaging were performed in accordance with a Temple University approved protocol. C57BL/6J and NSG mice (NOD.Cg-Prkdcscid Il2rgtm1Wjl/SzJ) were obtained from The Jackson Laboratory (C57BL/6J Stock No. 000664, NSG Stock No. 005557). [11]

### Atomic force microscopy of the human PDAC sections and force curves analysis

Tissue blocks were sectioned two days prior to the experiment. One section was designated for AFM, while adjacent sections were stained using H&E, P-c-Jun, and GFAP. Detailed protocols for tissue preparation and AFM procedures are provided in the supplementary methods.

### Atomic force microscopy on cultured cells to induce force and imaging to assess KTR distribution

Forces on nucleus and cytoplasmic parts of the cells were applied using a MFP-3D-BIO AFM microscope (Oxford Instruments) and cantilevers with 20 μm diameter borosilicate glass probes (CP-CONT-BSG-B, sQUBE). Cells were stimulated for 30-40 minutes and fluorescent images were acquired every 5 minutes with a 63x 1.4 NA oil immersion objective (Carl Zeiss).

### Model of murine pancreas fibrosis

Fibrosis in murine pancreas was achieved through intraperitoneal administration of cerulein. Ten to twelve week-old male and female mice received five intraperitoneal injections of cerulein (50 µg/kg) daily with an interval of one hour for 3 days per week for a total 4 weeks.

### Orthotopic transplantation of HEI-286-KTR SCs in murine pancreas

Orthotopic transplantation was performed as previously described.[18] Animals were anesthetized with isoflurane. Median laparotomy and pancreas exposure were performed in sterile conditions. HEI-286-KTR SCs expressing Clover fluorescence (200,000 cells in 10 µl PBS) were injected into the pancreas head under loop magnification. The abdomen was closed up with surgical sutures and pain medication was applied. Mice were followed for recovery and pain medication was administered daily for 72 hours.

### Intravital multiphoton microscopy

NSG mice were imaged 5 days post-injection with HEI-286-KTR SCs as previously described for other cell types.[19] Detailed description of the intravital multiphoton microscopy is provided in the supplementary methods.

### Live imaging in microchannels and compression in 6-well plate confiners

Microchannel dishes (4Dcell, Cat # MC004 (2-8 µm width), MC005 (10-20 µm width)) and 6-well cell confiner (4Dcell, Cat # CSOW 620) were utilized in accordance with the manufacturer’s protocol. Detailed descriptions of their use are provided in the supplementary methods.

### Quantification of c-Jun phosphorylation and nuclear rupture

The live c-Jun phosphorylation status in each cell was quantified using the KTR distribution by measuring the ratio of the cytosolic (C) to the nuclear (N) mean fluorescence intensity (C/N) for each cell using ImageJ and Prism softwares. Nuclear rupture was assessed by measuring NLS (nuclear localization signal) distribution. Bright NLS puncta observed before and after force application, likely reflecting peptide accumulation due to overexpression, were excluded from the analysis.

## RESULTS

### Stiffness of stromal tissue surrounding the nerves in PDAC correlates with P-c-Jun expression in SCs

To explore the effects of stiffness in PDAC with PNI, we used atomic force microscopy (AFM) to measure the stiffness of different areas of tumor (cancer cells, stroma, or nerves) in specimens collected from 13 different patients (**Figure 1 A-D**). Consistent with previous reports [20,21], areas with cancer cells had the lowest stiffness values (0.98 ±0.74 kPa) and stromal-rich areas had the highest stiffness values (3.74 ±2.79 kPa) reaching up to 14.01 kPa (**Figure 1D**). We also observed that nerve stiffness (1.79 ±1.24 kPa) was intermediate between stromal and cancer cell stiffness, but significantly higher in the presence of invading cancer cells (2.44 ±1.53 kPa vs 1.41 ±0.84 kPa). The stroma, which surrounds most of the nerves, had stiffness ranging from low to high values (0.43 to 14.01 kPa).

**Figure 1:**
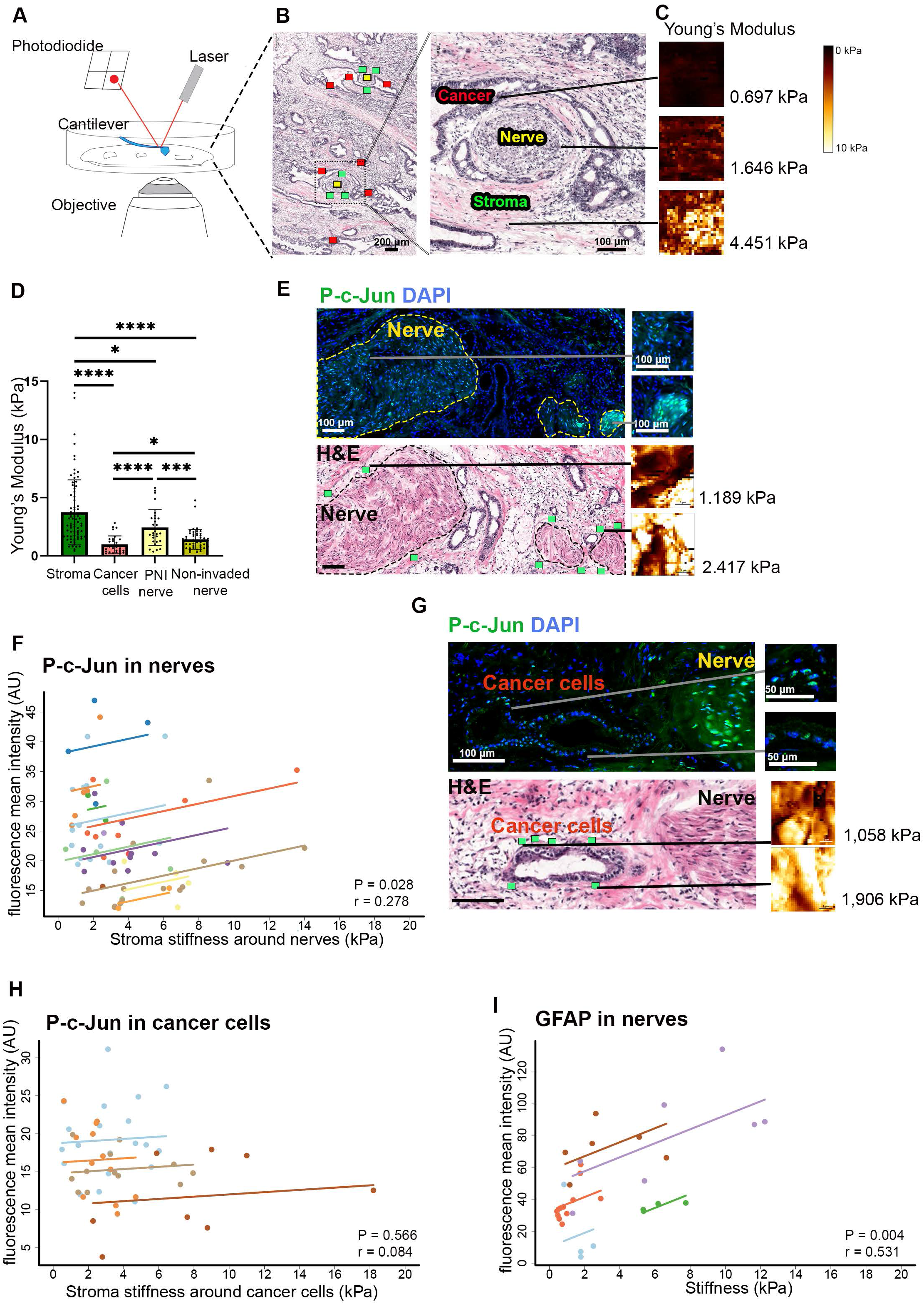
Tissue stiffness and SC activation in PDAC specimens. A: Atomic force microscopy setup to acquire elastic modulus in PDAC specimens. A sharp cantilever tip scans the surface, with deflections detected via laser reflection into a photodiode. B: H&E-stained sample showing nerves and PNI, with AFM measurement regions marked on an adjacent section. Colored boxes denote measured areas: red for cancer cells, yellow for nerve, and green for stroma. C: Young’s modulus and stiffness maps for cancer cells, nerve, and stroma, with values representing area-specific averages. D: Stiffness of the different PDAC components: stroma, cancer cells, PNI and non-invaded nerves obtained in samples from 13 patients. Student t test. *P≤0.05, *** P≤0.001, **** P≤0.0001 E: Different levels of P-c-Jun in nerves (dotted lines). P-c-Jun and H&E stainings on adjacent sections of a section used for AFM. Images on right shows low and high levels of nerve P-c-Jun (top) and stiffness maps with Young’s modulus mean values (bottom) around these nerves. Scale bar: 100μm. Green boxes indicate areas of AFM scanning. F: Correlation of P-c-Jun in nerves and corresponding stiffness of stroma surrounding these nerves using repeated measures correlation (rmcorr). The stiffness of stroma surrounding a nerve is the mean of 3 different measurements around the nerve. 13 patients. r= 0.278, P= 0.029. G: P-c-Jun in nerves and cancer cells. P-c-Jun and H&E stainings on adjacent sections of a section used for AFM. Images on right shows low and high levels of cancer cells P-c-Jun (top) and stiffness maps with Young’s modulus mean values (bottom) around these cancer cells. Scale bar 100μm. Green boxes indicate areas of AFM scanning. H: Cancer cell P-c-Jun and stiffness of stroma around the cancer cells do not correlate. (rmcorr. r= 0.084, P= 0.566) I: GFAP expression in single nerve fibers correlates with the stiffness of stoma around these fibers. (rmcorr, 5 patients, r = 0.531, P=0.004).

To test the hypothesis that mechanical stress on cells, as measured by tissue stiffness, activates SCs in the PDAC microenvironment, we measured stromal stiffness around nerves and assessed c-Jun phosphorylation status in these nerves using immunofluorescence on an adjacent section (**Figure 1E)**. We found a significant positive correlation between stromal stiffness and SC activation as measured by P-c-Jun staining in some individual patients (Patient 1 & 3) (**Supplementary Figure 1A**), and in the pool of the 13 patients using repeated measures correlation (rmcorr, r=0.278, p=0.028) (**Figure 1F)**. The correlation remained statistically significant if we removed Patient 1 and 3 form the cohort (r=0.382, p=0.011) (**Supplementary Figure 1B**). Notably, P-c-Jun in nerves did not correlate with nerve stiffness (**Supplementary Figure 1C**), likely because nerve tissue is intrinsically less stiff than the surrounding stroma. P-c-Jun was present in some cancer cells, but to a lesser extent than in nerves (**Figure 1G**), and P-c-Jun expression in cancer cells did not correlate with stromal stiffness measured around these cancer cells, suggesting a cell-specific mechanism of c-Jun kinase activation by the stiff stroma (**Figure 1H**).

We performed an orthogonal test of the correlation between stromal stiffness surrounding nerves and SC activation in areas with small nerve fibers (<10 um in diameter) in 5 patients. These fibers cannot be detected by H&E but express GFAP, another marker of activated SCs that is better suited to detecting single fibers because of its cytoplasmic localization. GFAP staining intensity in these fibers correlates with corresponding stromal stiffness in 3 individual patients and in a pool of 5 patients (**Figure 1I, Supplementary Figure 1D-E)**. These data suggest that stroma stiffness promotes SC activation via c-Jun phosphorylation, a sensitivity to mechanical stress that is not observed in cancer cells.

### Live monitoring of SC activation

To characterize SC activation, we monitored c-Jun phosphorylation in a human SC line, HEI-286, that expresses a c-Jun kinase translocation reporter (KTR) biosensor **(Supplementary Figure 2A-B)**. The KTR biosensor reports kinase activity by shuttling between the nucleus (N) and cytoplasm (C) based on phosphorylation of c-Jun residue after activation of a nuclear export signal and deactivation of a nuclear localization signal.[22,23] Anisomycin treatment increased P-c-Jun levels in HEI-286 KTR SCs as seen by Western-blotting (**Supplementary Figure 2C**), immunofluorescence of P-c-Jun (**Supplementary Figure 2D right**) and translocation of the KTR reporter (**Supplementary Figure 2D left**). Ratiometric expression of the sensor between cytoplasmic and nuclear compartments (C/N) provides a quantitative measure of c-Jun phosphorylation in each cell (**Supplementary Figure 2A-F. Movie 1**).

HEI-286 KTR SCs grown on a stiff substrate display a small but significant increase (10% of C/N ratio) in c-Jun activation (**Supplementary Figure 2G**), consistent with the observation that cell-surface integrins activate Jun kinase.[24] Treatment with the proinflammatory cytokine IL-6 and co-culture with pancreatic cancer cell lines MiaPaCa-2 and KPC induced a similar consistent 10% increase in c-Jun phosphorylation in HEI-286-KTR SCs compared to SC alone (**Supplementary Figure 2H-I**), consistent to previous reports.[11,12] Genes implicated in JNK activation, including integrins *ITGB3, ITGB5* as well as *PTK2, SRC* and *BCAR1*, which act on c-Jun kinase via focal adhesions [24], were more highly expressed in HEI-286 SCs co-cultured with MiaPaCa2 than HEI-286 SCs grown alone, supporting a mechanism of c-Jun kinase activation of SCs via cell-surface integrins (**Supplementary Figure 2J**). These *in vitro* experiments support a mild activation of SCs by pancreatic cancer cells, IL-6 and substrate stiffness.

### HEI-286-KTR SC activation occurs *in vivo* and depends on fibrosis

To further understand how stromal stiffness activates SCs *in vivo*, we monitored SC activation in normal and fibrotic murine pancreas after orthotopic HEI-286-KTR SC implantation **(Supplementary Figure 3A-C)**. After classifying the cells into three groups - activated, intermediate, and non-activated - we observed 3 times more activated SCs in fibrotic compared to untreated pancreas (**Figure 2A, Supplementary Figure 3D**). Interestingly, we observed a high level of cells in the intermediate group in normal pancreas, suggesting that differences between conditions may reflect variations in the degree of SC activation. We performed intravital imaging of SCs and surrounding collagen fibers using 4D multiphoton microscopy to investigate the switch from a non-activated to an activated state. Collagen fibers appeared wavy in normal tissue and straight in fibrotic tissue, as reported in cancer fibrosis (**Figure 2B**). [25,26] Most SCs were static (>90 %), while a few SCs moved within the surrounding extracellular matrix. These motile SCs squeezed through narrow areas and became activated, as detected by the redistribution of KTR in the cells, while changing their shape (**Figure 2C-D, Movie 2**). We also observed and confirmed by particle image velocimetry analysis a collagen displacement around and toward static activated cells (**Figure 2E-G, Movie 3**). Live imaging allowed us to observe a direct relationship between SC activation and mechanical constraints imposed by extracellular matrix features in the native pancreas environment.

**Figure 2.**
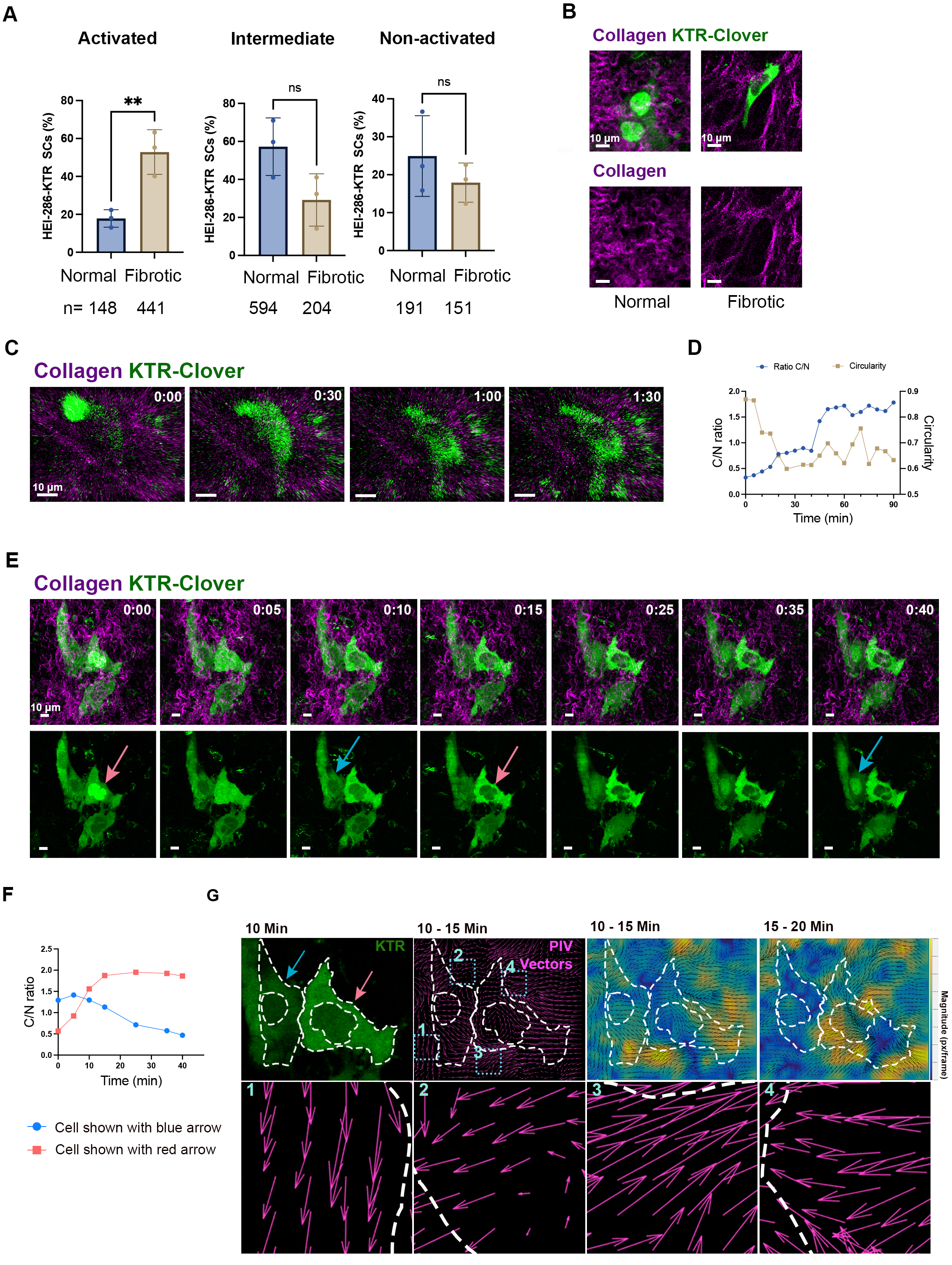
*In vivo* activation of HEI-286-KTR SCs in murine pancreas within a fibrotic microenvironment. A: Distribution of activated (C/N>1.2), intermediate (0.8<C/N <1.2), and non-activated (C/N<0.8) HEI-286-KTR SCs in normal and fibrotic pancreas (3 mice per condition with at least 100 cells per mouse). Student t test * p<0.05. B: Intravital multiphoton microscopy images of HEI-286-KTR SCs in normal and fibrotic murine pancreas showing representative non-activated SCs (C/N<1) in normal pancreas and an activated SC (C/N>1) in fibrotic pancreas. C: Still frames showing a HEI-286-KTR SC (**Movie S2**) migrating through extracellular matrix and undergoing activation when squeezing through narrow space. Time is h:min. Scale bar 10 µm. D: Analysis of C: Increase of C/N ratio and a concomitant decrease of circularity of the nucleus. E: Still frames showing HEI-286-KTR SCs (**Movie S3**) surrounded by moving collagen fibers. Red and blue arrows mark cells undergoing activation and deactivation respectively. Time is h:min. Scale bar 10 µm. F: Analysis of E: The cell marked with the red arrow shows an increased C/N ratio over time, indicating increase of P-c-Jun, whereas the cell marked with the blue arrow exhibits a decreased C/N ratio, consistent with decrease of P-c-Jun. G: HEI-286-KTR SCs and the corresponding graphical analysis of collagen fiber movements using particle image velocimetry. Spatial profiles of directionality and speed are depicted with vectors and a heat map for magnitude of the collagen movement occurring between 10 and 15 min, and between 15 and 20 min for 2 SCs, one getting activated (red arrow) and one getting inactivated (blue arrow). The shift of collagen fibers away from the inactivated and toward the activated cell is visualized by yellow and orange peaks of motion vectors. White dotted lines indicate nucleus and cell borders.

### Schwann cell activation depends on nuclear changes induced by mechanical forces

Static cells can be pushed by the stiff surrounding tissue and moving cells can meet resistance from solid components of the tissue. Such forces applied in a 3D environment can deform both plasma membranes and cell nuclei.[27] To test whether mechanical forces can induce c-Jun phosphorylation in SCs, we exerted differential forces on HEI-286-KTR SCs using three *in vitro* models: AFM (**Figure 3A-C**), microchannels of different width sizes (**Figure 3D-J**), and compression chambers (**Figure 3K-N**).

**Figure 3.**
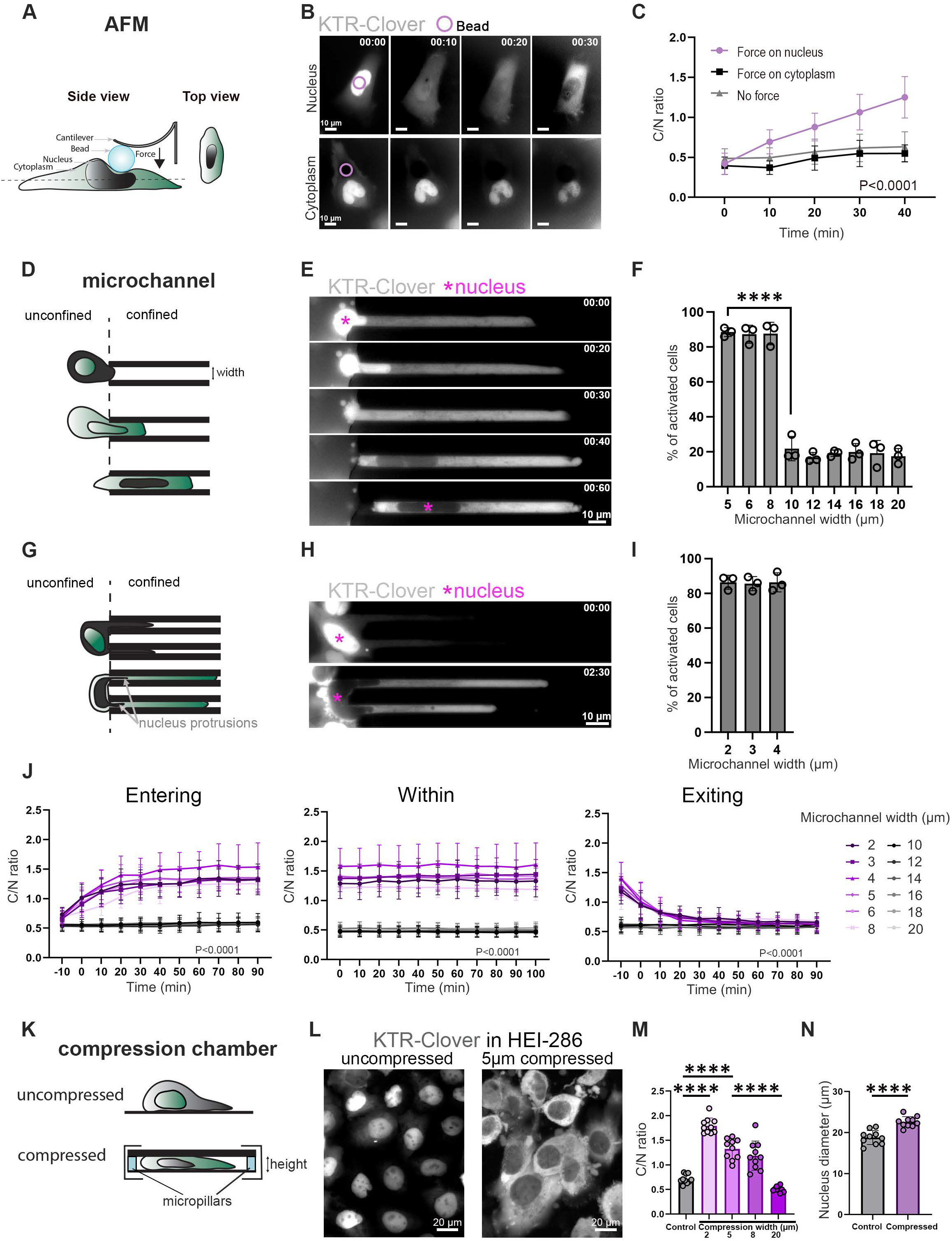
Mechanical forces increase c-Jun phosphorylation in HEI-286-KTR SCs. A: Schematic representation of AFM. A 20 μm bead is mounted on the cantilever tip to apply force either on or outside the nuclear region. B: c-Jun phosphorylation in live HEI-286-KTR cells stimulated with AFM. Upper row images represent a cell experiencing pressure in the nucleus region. Lower row images represent a cell experiencing pressure outside of the nucleus region. KTR translocates from a nuclear to cytosolic localization demonstrating c-Jun phosphorylation upon pressure on the nucleus. Circle indicates the position of the bead. Time is h:min. Scale bar: 10 μm. C: Quantification of c-Jun phosphorylation in HEI-286-KTR SCs in which AFM force is applied on the nucleus, outside of the nucleus or not applied. (n=10 cells per condition. P<0.0001, Repeated Measures ANOVA for condition and time effect, repeated twice). D: Schematic representation of an HEI-286-KTR SC navigating in microchannels (5 to 8 μm width). Cells are seeded in a chamber and enter microchannels of different width sizes in which they are confined. E: c-Jun phosphorylation of HEI-286-KTR SCs entering 5 μm width microchannel. At time 0, KTR fluorescence is mainly localized in the nucleus (asterisk). At time 60 min, the nucleus is within the microchannel and the fluorescence is mainly localized in the cytoplasm. Time: hour:min. Scale bar: 10 μm. F: Fraction of activated SCs (C/N>1) within microchannels of different widths (5 to 20 μm). (n>100 cells per microchannel width, 3 independent experiments). G: Schematic representation of an HEI-286-KTR SC in narrow microchannels (2 to 4 μm). Cells are seeded in the chamber and parts of them enter the microchannels. Nuclei of HEI-286-KTR SCs do not fully enter the narrow microchannels but form nucleus protrusions within the channels. H: c-Jun phosphorylation of HEI-286-KTR SCs partially inserted within a 4 μm width microchannel with nucleus protrusions. (asterisk is in the nucleus). Time: hour:min. Scale bar: 10 μm. I: Fraction of cells activated among the cells that are partially inserted within the narrow microchannels (2 to 4 μm width) (n> 100 cells per microchannel width, 3 independent experiments). J: c-Jun phosphorylation of HEI-286-KTR SCs entering, staying within and exiting microchannels (n=30 cells per microchannel). Timepoint 0 for entering and exiting cells corresponded to the first frame when the nuclei or parts of the nuclei were squeezed in the microchannels and when the nuclei were outside the microchannels, respectively. (P<0.0001, 2 way ANOVA, 5 μm vs 10 μm). K: Schematic representation of HEI-286-KTR SCs in compression chamber. HEI-286-KTR SCs seeded in glass bottom dish are compressed between two surfaces spaced by micropillars of different heights. L: c-Jun phosphorylation of uncompressed (left) and compressed (right) HEI-286-KTR SCs. Compression height 5 μm. Scale bar 20 μm. M: Effect of compression on c-Jun phosphorylation in HEI-286-KTR SCs. (n=10 cells per condition, mean ± SD, two ways Student t-test, repeated 3 times) N: Effect of compression on nucleus diameter in uncompressed and compressed HEI-286-KTR SCs (5 μm). (n=10 cells per condition, mean ± SD, two ways Student t-test, repeated 3 times).

Using AFM, we exerted forces focally on either nuclear or cytoplasmic areas of the cells and monitored KTR distribution within each cell. We also monitored nuclear rupture using the nuclear marker NLS (**Supplementary Figure 4A**). Applying a force to nuclei resulted in a strong increase of c-Jun phosphorylation (C/N > 1) - without rupturing the nuclear membrane - compared to control non-stimulated cells, and compared to cells in which forces were applied in cytoplasmic regions (**Figure 3B-C**). This suggested that forces applied on SC activate SC and the force-sensing mechanism inducing c-Jun phosphorylation was localized in the nuclear region of the SCs.

We next monitored c-Jun phosphorylation in HEI-286-KTR SCs entering polydimethylsiloxane microchannels by time-lapse imaging. Cells were placed into unconfined 2-dimensional chambers and subsequently migrated to 3D-confined microchannels of 2 to 20 μm width (**Figure 3D-J and Movie 4**). SCs fully entered microchannels with a width of 5 μm and above, while only portions of the SC cytoplasm and nuclear protrusions entered microchannels with a width of 2, 3 and 4 μm. The observation of compressed nuclei that do not enter narrow microchannels is consistent with studies reporting that the nucleus is the most rigid organelle of the cell [28]. Within the 10-μm wide and larger microchannels, only 20% of cells showed evidence of activation (C/N>1), whereas 80% of cells were activated in the narrower microchannels (5 to 8 μm width), suggesting that confinement induced SC activation (**Figure 3E-F**). Cells attempting to enter the narrow microchannels (2 to 4 μm) and undergoing nuclear compression also showed c-Jun activation (**Figure 3H-I**). This was reminiscent of squeezing cells within the fibrotic environment seen *in vivo* (**Figure 2**). Interestingly, c-Jun phosphorylation remained high when cells stayed within narrow microchannels, and stayed low within large microchannels (**Figure 3J**). Furthermore, c-Jun phosphorylation increased with SC entry into the microchannels and decreased upon SC exit, indicating that activation is reversible (**Figure 3J, Supplementary Figure 4B**). We previously showed that HEI-286 SCs grown in 3D Matrigel matrix organize longitudinally in stellate structures[10,11] with widths of 10 μm (9.65 ± 0.65, n=10), which corresponds to the microchannel width beyond which cells are not activated. This suggests that SCs create an environment in which c-Jun is not phosphorylated.

We next placed HEI-286-KTR SC in compression chambers at predefined heights (2, 5, 8 and 20 μm) to apply direct force (**Figure 3K-N**), and monitored nuclear size and nuclear envelope rupture by tracking the localization of a fluorescent nuclear localization signal (NLS) peptide.[29] Nuclear size increased in compressed cells (**Figure 3N**), but the NLS signal stayed inside the nucleus of cells undergoing 5-μm and 8-μm compression, suggesting that the stretched nuclear membranes remained intact (**Supplementary Figure 4C**). However, NLS became largely cytotosolic in cells compressed to 2 μm, indicating nuclear rupture (**Supplementary Figure 4C**). SCs compressed to 5 and 8 μm showed higher c-Jun phosphorylation than uncompressed SCs and SCs compressed to 20 μm (**Figure 3M)**. The increase of P-c-Jun in compressed cells was also confirmed by Western-blotting and Immunofluorescence (**Supplementary Figure 4D-E**). GFAP, another marker of SC activation was also increased in compressed cells (**Supplemenatry Figure 4F**). These results demonstrate that c-Jun is phosphorylated in SCs under mechanical compression leading to stretching of the nuclear membrane. Collectively, our three different applications of force support that mechanical compression acts through nuclear membrane distortion to induce P-c-Jun.

### Expression profiles of compressed SCs reveal an AP1-dependent stress response

We sought to learn how gene expression changes in response to short and long-term compression. We thus profiled HEI-286 SCs with RNA-seq after a brief 5-min compression, and after 4 h, a time period corresponding to previous studies of cell compression in other cell types[30] We observed a striking increase in the expression of genes encoding the AP-1 transcription factor complex or its interaction partners, including *FOS, FOSB, NFATC2, EGR1* and *ATF3* both 5 min and 4 h after compression (**Figure 4A-B**). This is consistent with our initial observations of increased c-Jun phosphorylation in compressed cells (**Figure 3)** since P-c-Jun dimerizes with FOS to form the AP-1 complex. Pathway analysis revealed ‘AP-1 transcription factor network’ as the most significantly upregulated pathway in the NCI Nature database at both timepoints, and for the genes that increase progressively between 5 min and 4 h (**Figure 4C, 4E)**. Scoring gene sets confirms the implication of the upregulated genes in the ‘AP-1 transcription factor network’ in compressed as compared to uncompressed cells (**Figure 4D**), which include *FOS, FOSB, ATF3, EGR1, CSF2, NFATC2, JUN, JUNB, JUND, MT2A, CYR61* and *COPS5* (**Figure 4F**).

**Figure 4.**
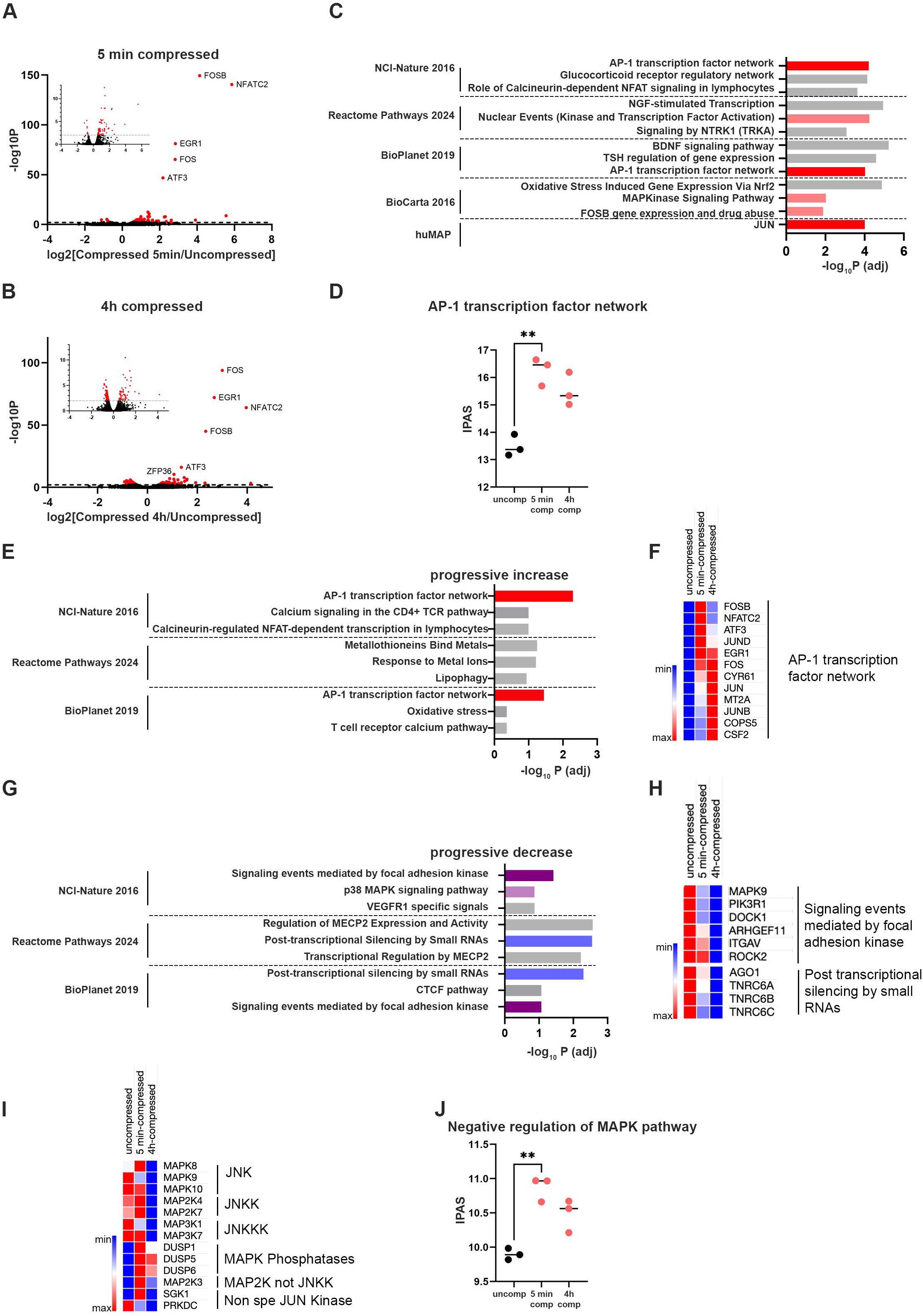
Transcriptomic analysis of compressed HEI-286 SCs. A and B: Volcano plots showing differentially expressed genes between uncompressed HEI-286 SCs and compressed HEI-286 SCs for 5 min (A) and 4 hours (B) (5 μm height compression). C: Top three enriched pathways of upregulated genes in 5 min-compressed HEI-286 SCs in indicated data sets. D: Inferred pathway activation/suppression (IPAS) scores for ‘AP-1 transcription factor network’ pathway in uncompressed and compressed HEI-286 SCs. E: Top three enriched pathways of genes that gradually increase expression from uncompressed state to 5 min and to 4 h compressed state in indicated data sets. F: Heatmap of genes of the ‘AP-1 transcription factor network’ pathway that are upregulated after 5 min compression and/or 4h compression, ordered by appearance time. G: Top three enriched pathways of genes that gradually decrease expression from uncompressed state to 5 min and to 4 h compressed state in indicated data sets. H: Heatmap of the downregulated genes that gradually decrease expression from uncompressed state to 5 min and to 4 h compressed state from the ‘focal adhesion kinase’ and ‘post transcriptional silencing by small RNAs’ pathways. I: Heatmap of expression of genes affecting c-Jun kinase signaling pathway including specific and non-specific Jun kinases, Jun kinase kinases, and Jun kinase phosphatases (DUSP). J: IPAS scores for genes set corresponding to ‘Negative regulation of MAPK pathway’ (Reactome).

Interestingly, compression also increased expression of immune-related pathways, including ‘Role of calcineurin-dependent NFAT signaling in lymphocyte’, ‘T cell receptor calcium pathway’, ‘calcium signaling in the CD4+ TCR pathway’ **(Figure 4C, 4E)**. These results suggest that mechanical stress may also have direct and/or indirect immunomodulatory effects on the pancreatic stroma

### Mechanical force induces a negative regulation of pathways related to JNK

Surprisingly, genes of the ‘signaling events mediated by focal adhesion kinase’ and ‘p38 MAPK signaling’ pathways, which are related to JNK signaling, showed progressive downregulation during compression. They included *MAPK9* (or JNK2), *PIK3R1, ARHGEF11, DOCK1, ITGAV* and *ROCK2* (**Figure 4G-H**). Other JNK-related genes such as *MAPK8* (*JNK1*) and *MAPK10* (JNK3), as well as the members of JNK kinase (JNKK) *MAP2K4* and *MAP2K7*, and JNKK kinases *MAP3K1* and *MAP3K7* also showed a decrease of expression during compression (**Figure 4I)**. We also noted an increased expression of genes encoding the phosphatases DUSP1, DUSP5 and DUSP6, which are negative regulators of MAPK and JNK activity, in compressed cells as compared to uncompressed cells (**Figure 4I)**. IPAS scores confirmed the negative regulation of MAPK pathway genes in compressed cells (**Figure 4J**).

These data led us to hypothesize that compression-induced c-Jun phosphorylation leads to negative feedback to suppress conventional JNK activity, and that c-Jun phosphorylation in SCs in this context is mediated through a non-canonical kinase interaction. Serum/glucocorticoid regulated kinase 1 (*SGK1*) may be one such non-canonical mediator, since *SGK1* was an upregulated kinase in our model and was shown to activate the AP-1 complex in breast cancer cells[31] These data collectively show that the compression of HEI-286 SCs leads to c-Jun phosphorylation, an increase of AP-1 formation, and surprisingly transcriptomic changes indicative of negative regulation of JNK activation and canonical downstream signaling, suggesting an alternate mechanism of c-Jun phosphorylation.

### cPLA2 is a nuclear mechanosensor involved in SC activation

Several of our observations suggest that the sensing mechanism for force-induced c-Jun phosphorylation is in the nucleus. We observed that c-Jun phosphorylation occurs specifically when force is applied (1) to the nucleus but not the cytoplasm using AFM (**Figure 3B and 3C**), (2) in synchrony with nuclear membrane distortion in the microchannel assay and *in vivo* experiments (**Figure 3E-J and Figure 2C-D**), and (3) to deform nuclei in SCs confined the compression chambers (**Figure 3L-N**). Therefore, we hypothesized that sensors responsible for c-Jun phosphorylation upon mechanical forces are present at the nucleus.

The calcium-dependent cytosolic phospholipase A2 (cPLA2) is a mechanosensor that transits from the nucleoplasm to the inner nuclear membrane when the nuclear envelope experiences tension.[32–34] We expressed cPLA2-mKate in HEI-286-KTR SCs and applied force to the nuclei using AFM. Imaging revealed that cPLA2-mKate translocated from the nucleoplasm to the nuclear envelope within one minute; subsequently, KTR in the same cells translocated from the nucleus to the cytoplasm at a slower rate (**Figure 5A**). We hypothesized that the mechanism of c-Jun phosphorylation in compressed SCs depends on cPLA2. We placed HEI-286-KTR SCs into 5-μm high chambers and compared cells treated with vehicle control, specific JNK inhibitor SP600125, a non-specific JNK (SGK1) inhibitor GSK 650394 or a cPLA2 inhibitor pyrrophenone. As anticipated, we observed c-Jun phosphorylation in the control compressed SCs; however, this was abrogated by all three of the pharmacological treatments (**Figure 5B-C**). We also observed a significant inhibition of c-Jun phosphorylation in compressed HEI-286-KTR SCs that were genetically ablated for cPLA2 using CRISPR-Cas9 (**Figure 5D-E, Supplementary Figure 5A**). In addition, P-c-Jun and GFAP immunofluorescence stainings on compressed HEI-286-KTR SCs were lower in cPLA2 KO cells than in control cells **(Figure 5F)**. Importantly, treatment with arachidonic acid, a metabolite product of cPLA2 activation, induced c-Jun phosphorylation on HEI-286-KTR SCs (**Figure 5G)**. Furthermore, c-Jun phosphorylation was lower in cPLA2 KO HEI-286-KTR cells injected in murine fibrotic pancreata than in control HEI-286-KTR SCs (**Figure 5H**).

**Figure 5.**
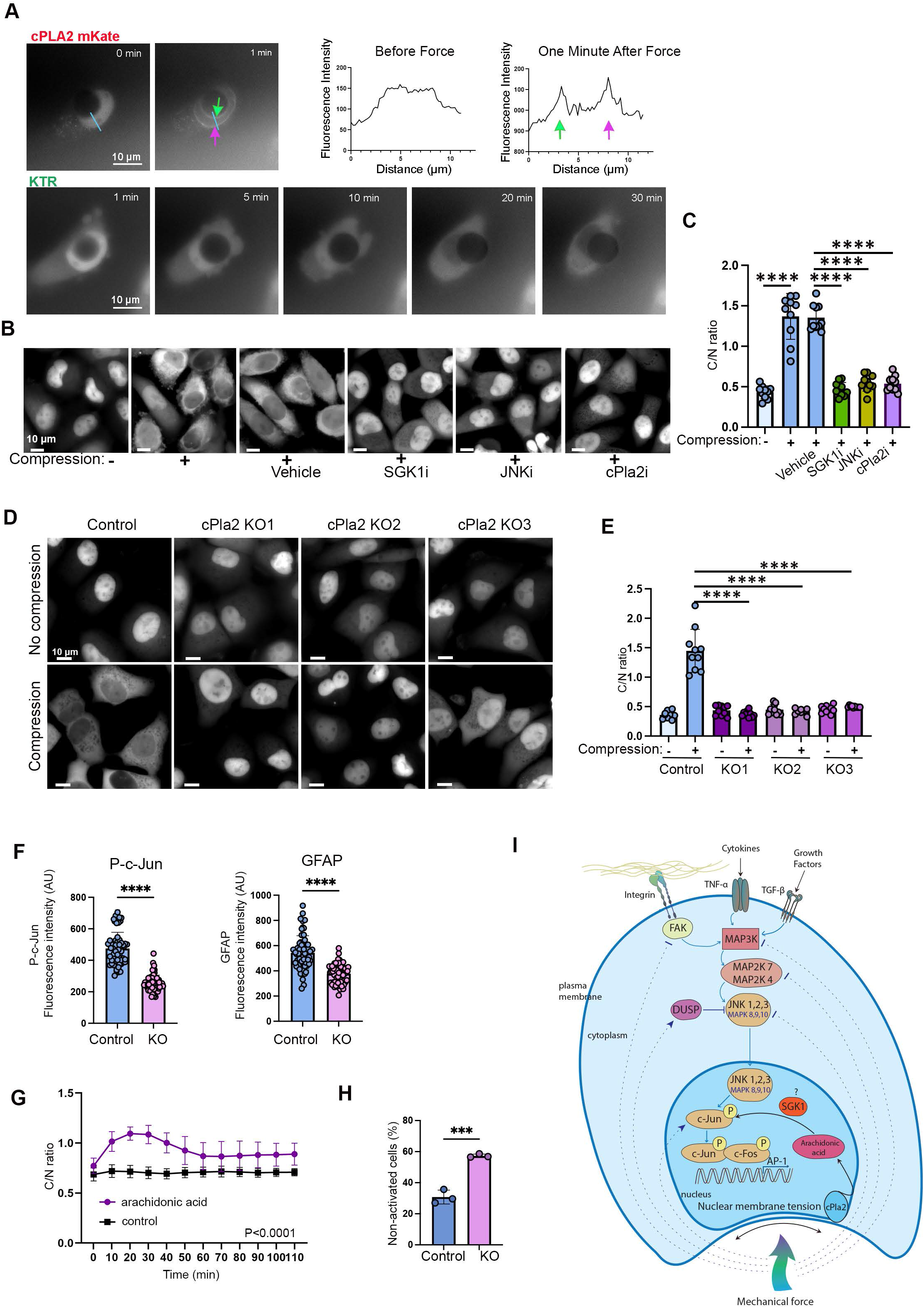
Molecular mechanisms of c-Jun phosphorylation in SCs induced by mechanical forces. A: cPLA2 translocation from nucleus to nuclear membrane and KTR translocation from nucleus to cytoplasm in a HEI-286-KTR cPLA2 mKate cell after force was applied to the nucleus using an AFM cantilever (top). Quantification of cPLA2-mKate intensity along the blue line drawned in the images. Colored arrows indicate the nuclear membranes (Note: differences in Y-axis values are due to changes in exposure time between the two images to compensate photo-bleaching). Scale bar 10 μm. Repeated 6 times. B: Effect of JNK inhibitor (SP600125), cPla2 inhibitor and SGK1 inhibitor on c-Jun phosphorylation in compressed HEI-286-KTR SCs. Concentration of all inhibitors 10 μM. Compression height 5 μm. Scale bar 10 μm. C: Quantification of A. (n=10 cells per condition, mean ± SD, two-way Student t-test, ****p<0.0001, repeated 3 times) D: Effect of cPLA2 depletion on c-Jun phosphorylation in compressed HEI-286-KTR SCs. Control are cells expressing a non-targeting control gRNA, cPla2-KO1, -KO2 and -KO3 are cPla2 knock-out cells expressing 3 different sgRNA guides targeting cPla2. Compression height 5 μm. Scale bar 10 μm. E: Quantification of C. (n=10 cells per condition, mean ± SD, two-way Student t-test, ****p<0.0001, repeated 3 times). F: P-c-Jun and GFAP immunostaining in compressed in control and cPLA2 KO HEI-286-KTR SCs (n>50 cells per condition, mean ± SD, two-way Student t-test, ****p<0.0001, repeated 3 times). G: c-Jun phosphorylation in HEI-286-KTR cells treated with arachidonic acid (1 μM). (n=10 cells per condition, mean ± SD, two-way ANOVA, ****p<0.0001, repeated 3 times) H: Quantification of non-activated SCs (C/N<0.8) among control and cPLA2 KO HEI-286 KTR SCs injected in fibrotic pancreata (n=3 sections, 15 to 44 cells/section, sections distanced by 75 μm, mean ± SD, two-way Student t-test, ***p<0.001, one repeat) I: Schematic representation of mechanisms of c-Jun phosphorylation in SCs showing a separate signaling mechanism induced by mechanical force. In addition to the c-Jun signaling pathway stimulated by integrin, cytokine and growth factors that is transmitted via plasma membrane receptors and implicates FAK, MAP3K, MAP2K and MAPK/JNK (scheme upper part), a mechanism of c-Jun phosphorylation is activated by mechanical force inducing nuclear membrane tension, cPLA2 activation and arachidonic acid release (scheme lower part). Mechanical forces also inhibit the plasma membrane sensing and signaling mechanism by decreasing expression of FAK, JNKs and JNK Kinase and increasing DUSP expression.

Together, these data support cPLA2 as a nuclear mechanosensor in SCs that induces c-Jun phosphorylation. We propose a model in which mechanical forces that deform the nuclear membrane activate cPLA2, which induces AP1 complex formation and releases arachidonic acid; this activates a non-specific c-Jun kinase to phosphorylate c-Jun, while suppressing canonical c-Jun activation through MAPK and FAK activity (**Figure 5I**).

### Differential gene expression response in SCs and pancreatic cancer cells

The response to stress is one of the most common hallmarks of tumors[35] We compared upregulated genes of compressed HEI-286 SCs to that of pancreatic cancer cells MiaPaCa-2 and Panc01 under similar compression conditions. Cancer cells upregulated genes of the AP-1 transcription factor network, but fewer genes from the pathway were differentially expressed and the differentially expressed genes had lower fold changes (**Figure 6A-B, Supplementary Figure 5B**). Genes that were significantly upregulated in compressed HEI-286 cells but not in MiaPaCa-2 or Panc01 included *EGR1, CSF2, DUSP1, NFATC1, ETS1, HIF1A, FOSL1, MT2A, IL6, PLAU* and *JUNB*. These data showed an activation of AP-1 in all cells in response to compression, but the response was far more pronounced in HEI-286 SCs as compared to MiaPaCa-2 or Panc01 cancer cells.

**Figure 6.**
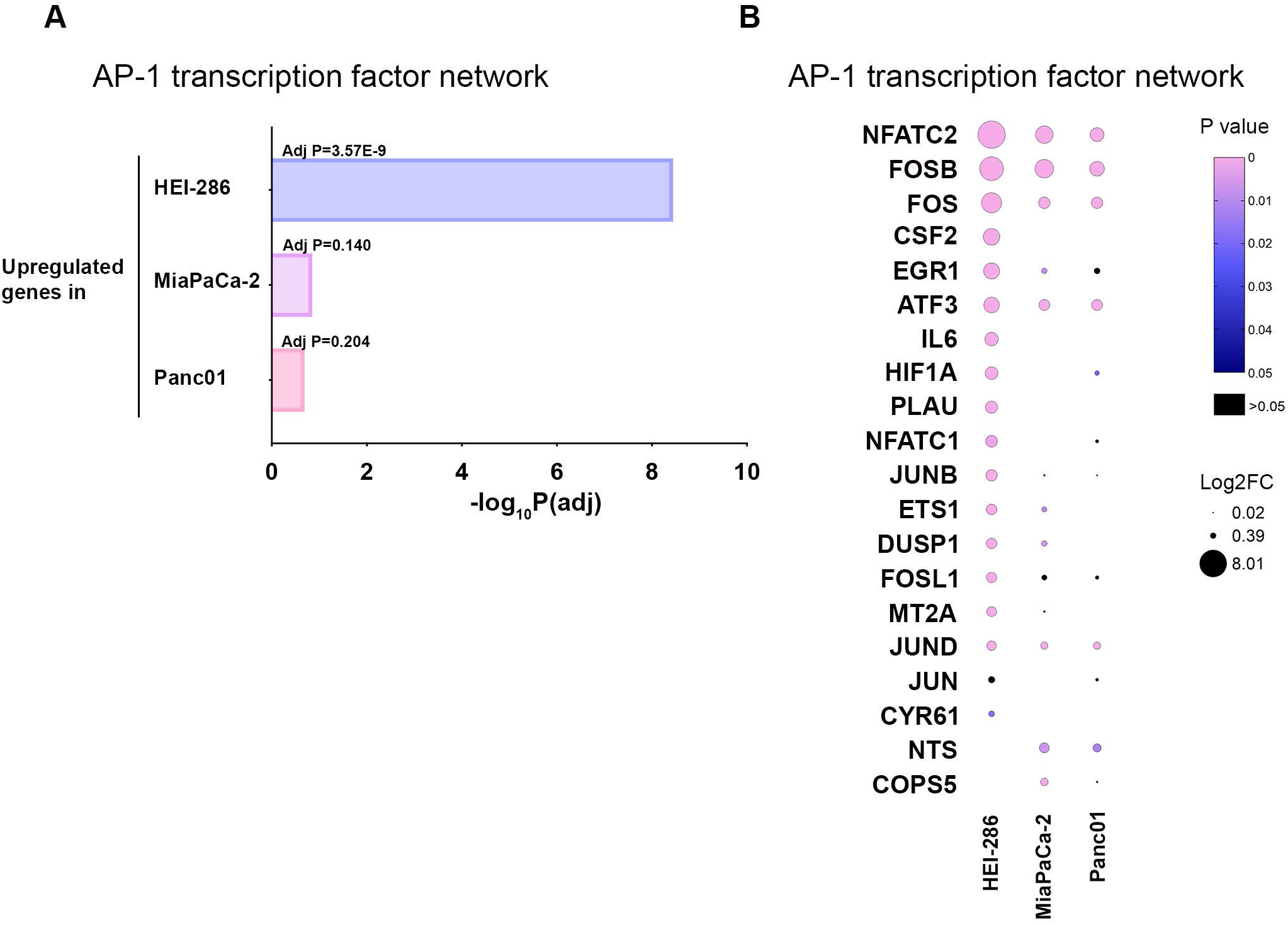
SC and cancer cell response to mechanical forces. A: Degree to which the ‘AP-1 transcription factor network’ pathway is represented in the compressed upregulated genes from HEI-286, MiaPaCa-2 and Panc01. B: ‘AP-1 transcription factor network’ genes upregulated in HEI-286 SCs, MiaPaCa-2 and Panc01 after compression. Missing dots indicate absence of gene expression change or downregulation of gene expression (NCI-Nature 2016 data set)

## DISCUSSION

Understanding how TME components regulating PDAC malignancy affect each other is key to unravel the full complexity of PDAC biology. Activated SCs have emerged as a significant protumorigenic player in PDAC.[10,11,17] In this study, we show how fibrosis impacts the activation of SCs. The fibrotic stroma of PDAC creates a stiff tissue environment that exerts resistance forces on SCs, which respond by activating the AP-1 transcription factor complex via phosphorylation of c-Jun in a manner that depends on nucleus deformation. This AP-1 response to mechanical force in SCs is greater than cancer cell response to forces.

We previously showed that SCs become activated and reprogrammed in cancer [10,11], and our results now elucidate a mechanism of activation. SC reprogramming depends on c-Jun and allows SC to facilitate cancer cell spreading.[11] In this study, we showed that fibrosis creates a more favorable environment to induce c-Jun activation in SCs that sense mechanical forces. The live imaging experiments enabled us to directly observe the relationship between compression of cells and cell nuclei (indicating response to mechanical forces in the environment) and SC activation (which immediately followed compression). This finding presents a way in which fibrosis contributes to PDAC malignancy.[2,36] Because activated SCs are present before the onset of PDAC[15], at a time when fibrosis is present in the pancreas, it is likely that SCs are also activated by fibrosis at this premalignant stage. SCs are also reprogrammed in injured nerves in a c-Jun dependent manner during wound repair[37], a process that involves deposition of excess extracellular matrix, similar to fibrosis.[38]

We applied a 120 nN force in our AFM experiments to induce c-Jun phosphorylation, a magnitude that lies within the physiological range of cellular-scale mechanics. While this value exceeds typical molecular forces in the picoNewton range such as forces generated by motor proteins or protein unfolding, cellular forces commonly reach tens to hundreds of nanonewtons, including cell traction forces and cancer cell indentation during invasion. [39–42] In this context, our observation that forces below 120 nN do not induce KTR translocation in HEI-286 cells suggests that c-Jun activation in SCs is not driven by molecular-scale forces but rather by higher-magnitude, cell-generated mechanical stresses. This supports a model in which global cellular forces capable of deforming the nucleus are required to trigger mechanosensitive signaling.

These data challenge the traditional view of c-Jun activation via a signaling cascade triggered by a stimulus detected at the plasma membrane. Stimuli triggering JNK include growth factors, proinflammatory cytokines (TNFα, IL-6) and UV radiation stress. Extracellular matrix components in the stroma can also stimulate JNK via integrin binding, which activates FAK and then Jun kinase.[24] JNK is activated by these stimuli after sequential protein phosphorylation through the MAP kinase module, in which MAP3Ks (JNKKK) phosphorylates and activates MAP2Ks (JNKK). In contrast, our data are consistent with a phosphorylation of c-Jun in SCs triggered by mechanical forces applied on nuclei and that inhibit expression of JNKK and JNKKK. We propose that under force, stretching of the nuclear membrane activates cPLA2 and releases arachidonic acid, which then induces c-Jun phosphorylation independently of JNK. It is noteworthy that cPLA2 mRNA expression is higher in human pancreatic cancer tissue than in normal pancreatic tissues.[43]

A candidate kinase for c-Jun phosphorylation through this mechanism is SGK1, a non-specific JNK whose gene expression increased upon SC compression in our study (**Figure 4**). We propose that both plasma and nuclear membrane detection can induce c-Jun phosphorylation in SCs. The contributions of membrane detection can vary among different cell types. The AP-1 response to mechanical forces is greater in SCs than in the cancer cells, as revealed by transcriptome analysis of compressed HEI-286 SCs and pancreatic cancer cells, MiaPaCa-2 and Panc01 (**Figure 6**) and by comparing the correlation between the stiffness of the stroma surrounding nerves and cancer cells with P-c-Jun content in these cells in PDAC specimens (**Figure 1**). It is noteworthy that the compressed gene signature is a stress response gene signature, sharing similarities with the stress-response program described in PDAC and other cancer types.[35,44] Stress-like cancer cell states have been detected from early tumorigenesis, and are associated with higher tumor-seeding capabilities and drug-resistance properties.[44] These genes constitute the second most abundant gene program among 41 consensus meta programs derived from 1163 samples covering 24 cancer types.[35]

In addition to add to the growing appreciation of the role of SCs in cancer malignancy, we report here that SCs are remarkable sensors for mechanical forces in pancreatic cancer. This allows the initiation of their reprogramming, which is essential for several SC functions, including the secretion of factors that promote cancer invasion[45], the organization of tracks for cancer[11] and the support of neurogenesis in cancer.[16]

A key limitation of this study is the use of human SCs injected into the murine pancreas, which may not fully recapitulate the behavior of endogenous SC. Differences in interactions with neurons, extracellular matrix, and immune cells could influence the activation state and mechanosensitive responses of the SCs. Therefore, while our findings demonstrate that SCs can respond to the fibrotic pancreatic microenvironment and exhibit mechanosensitive signaling *in vivo*, these results should be interpreted as a proof of concept rather than a representation of resident SC behavior. Future studies using genetically engineered mouse models with knock-in alleles will be important to validate these observations in endogenous SC populations.

Our findings reveal a critical role for fibrosis with associated mechanical forces in inducing SC activation in pancreatic cancer and could provide clues for developing new therapies to attenuate cancer dissemination by regulating SC activation.

## ETHICS STATEMENT

### Patient consent for publication

Not applicable.

### Ethics approval

This study involves human participants and was approved by the Institutional Review Board at Memorial sloan Kettering Cancer Cenetr (IRB-15052). All animal experiments were performed in accordance with the Memorial Sloan Kettering Cancer Center Institutional Care and Use of Animals Committee (IACUC)-approved protocols.

## ACKNOWLEDGEMENTS

We acknowledge the technical services provided by MSK core facilities which include the Molecular Cytology, the Gene Editing and Screening, the Flow Cytometry, the Integrated Genomics Operation, the Bioinformatics, and the Animal Imaging core facilities, funded by CCSG P30-CA08748. We thank Nada Wahman and Kerisha Williams for assistance with AFM experiments, and Ning Fan for assistance with the preparation of histological sections.

We also thank Dr. Niethammer for providing the cPLA2-mKate plasmid. Funding was provided by the Deutsche Forschungsgemeinschaft (DFG, German Research Foundation) 491318253 (PS), Hirshberg Foundation Seed Grant (SD), NIH R01CA219534 (RW), NIH NCI R01 CA230777 (BG); American Cancer Society 134415-RSG-20-34-01-CSM (BG), DOD BCRP

BC230197 (BG), PA-CURE (BG), WW Smith Charitable Trust (BG), METAvivor (IVQ). We thank Julia Simundza and Tal Nawy for critical reading and editing of the manuscript.

## FOOTNOTES

### Patient and public involvement

Patients and/or the public were not involved in the design, or conduct, or reporting, or dissemination plans of this research.

