## Supplemental Materials for "Pancreatic cancer fibrosis activates protumorigenic Schwann cells through a nuclear mechanosensing mechanism"

### SUPPLEMENTARY METHODS

#### Cell lines and cell culture

The human non-neoplastic HEI-286 SC line, the pancreatic cancer cells MiaPaCa-2-RFP and Panc01-RFP, the murine pancreatic cancer cell line Panc02 and KPC were previously described.[1] The murine fibroblast cell line NIH 3T3 was purchased from ATCC (Stock No. CRL-1658) and previously described.[2] HEI-286-KTR expressing Clover fluorescence was generated using pENTR-JNKKTR Clover (Addgene plasmid # 59139) vector. HEI-286-KTR pRFP-NLS was generated using pRFP-NLS (Addgene plasmid #105296) vector. HEI-286-KTR cPLA2mKate was generated using cPLA2-mKate plasmid (Addgene plasmid # 163844).

PLA2G4A KO HEI-286-KTR cell line was generated using CRISPR–Cas9 technology. Three guides were used for each gene. Constructs were made at the MSK Gene Editing and Screening Core Facility using pSpCas9(BB)-2A-Puro (PX459) V2.0 vector (Addgene plasmid #62988) and the following targeting oligonucleotide sequences:

| | |
|---|---|
| *PLA2G4A*_1 F | CACCGAAGAGAGGCAAAGGACACC |
| *PLA2G4A*_1 R | AAACGGTGTCCTTTGCCTCTCTTC |
| *PLA2G4A*_2 F | CACCGAACAAAGTGGATAATGGTT |
| *PLA2G4A*_2 R | AAACAACCATTATCCACTTTGTTC |
| *PLA2G4A*_3 F | CACCGAAGAATCAAGGGATATGGC |
| *PLA2G4A*_3 R | AAACGCCATATCCCTTGATTCTTC |

DNA constructs were nucleofected in HEI-286 cells using Amaxa nucleofector (program T-020). A control cell line was generated using a non-targeting sequence. Stable monoclonal cell lines were generated after selection with puromycin (10 µg/mL). Depletions of PLA2G4A in PLA2G4A KO HEI-286-KTR cells were verified by qRT-PCR using standard protocols.

All cells were cultured in 5% $CO_2$ at 37°C in Dulbecco's modified Eagle's medium (DMEM, Gibco) containing 10% FBS (Gemini) and 50 U/mL penicillin/streptomycin (Gibco). The culture medium for PLA2G4A KO HEI-286-KTR cells was supplemented with puromycin (10 µg/mL). The culture medium for cells expressing GFP and RFP was supplemented with G418 (50 μg/mL). Cell lines were routinely screened to avoid Mycoplasma contamination and maintained in a humidified chamber with 5% $CO_2$ at 37°C.

**Antibodies and reagents**

Antibodies include anti-GFAP (BioLegend, previously Covance, PRB-571C), anti-P-c-Jun (phospho S63) (abcam, ab32385), anti-c-Jun ([E254], abcam, ab32137), anti-S100 (Sigma-Aldrich, MAB079-1), anti-cytokeratin (BioLegend, 628601), anti-vinculin (VIN-11-5, Sigma Aldrich, V4505), anti-SOX10 ([5H7L26], Thermo Fisher Scientific, Cat#: 703439), anti-EGR2 (Novus Biologicals, Cat#: NB110-59723T), anti-GAP43 ([EP890Y], abcam, ab75810).

Chemicals include cPLA2α Inhibitor II, Pyrrophenone (10 μM) (Sigma-Aldrich, 530538), SP600125 (10 μM) (Tocris, Cat. No. 1496), SGK1 inhibitor, GSK 650394 (10 μM) (Tocris, Cat. No. 3572), Arachidonic acid (1 μM) (Sigma-Aldrich, A3611), IL-6 (100 ng/ml) (PeproTech, 200-06), Anisomycin (100 ng/ml) (Tocris, Cat#1290).

Plasmids obtained from Addgene were: pENTR-JNKKTRClover (Addgene plasmid # 59139); pRFP-NLS (Addgene plasmid #105296); pSpCas9(BB)-2A-Puro (PX459) V2.0 (Addgene plasmid #62988). Constructs for PLA2G4A KO were made using CRISPR-Cas9 technology at the MSK Gene Editing and Screening Core Facility. cPLA2-mKate plasmid (Addgene # 163844) was from Philipp Niethammer's laboratory.

**Immunofluorescence staining**

Frozen sections from human and mouse specimens were fixed with 4% paraformaldehyde. After permeabilization and blocking in 3% horse serum with 0.1% Triton X-100/PBST for 1 hour, sections were incubated with primary antibodies (anti-P-c-Jun 1:1000, anti-GFAP 1:5000, anti-SOX10 1:1000, anti-EGR2 1:1000 or anti-GAP43 1:1000) diluted in 0.1% horse serum and 0.1% Triton X-100/PBS overnight at 4 °C. The detection was performed using an appropriate fluorescent secondary antibody (Alexa Fluor 488, 568). Slides were mounted in DAPI containing antifade medium and scanned using Mirax Scanner (Zeiss), sections were analyzed using ImageJ.

HEI-286-KTR SCs were immuno-stained with anti P-c-Jun and anti-GFAP antibodies after cells were compressed or not for 10 min using the 6-well cell confiner (4Dcell, Cat # CSOW 620) with the 5 μm confinement slides. After compression, cells were immediately fixed with 4% paraformaldehyde, blocked with 3% horse serum with 0.1% Triton X-100/PBST for 1 hour, incubated with primary antibodies (anti-P-c-Jun 1:1000, anti-GFAP 1:5000) overnight at 4 °C and incubated with secondary antibodies (Alexa Fluor 568) for 1h. Cells were imaged for P-c-Jun or GFAP (Alexa 568) and KTR (Clover) using an inverted Zeiss Axio Observer microscope with an EC Plan-Neofluar 40X 0.3NAPh1 lens 40x objective and analyzed using Zen Software (Carl Zeiss) and Fiji.

**Western blot**

Cells were lysed using Pierce IP Lysis Buffer (Thermo Scientific, Cat# 87787) containing Protease/Phosphatase Inhibitor Cocktail (Cell Signalling, Cat# 5872) on ice for 30 minutes. Lysates with 30μg of protein were added to Blue Protein Loading Dye (BioLabs, Cat# B7703S) and Reducing Agent (BioLabs, Cat# B7705S) and incubated at 98°C for 8 minutes. Samples were separated by SDS-PAGE (4-15%) and transferred onto a nitrocellulose membrane. The membrane was blocked using Intercept Blocking Buffer (LI-COR, Cat# 927-70001) for one hour. The membrane was then incubated with primary antibodies diluted in blocking buffer overnight at 4°C. On the next day, the membrane was incubated with fluorescent-conjugated secondary antibodies (LI-COR, IRDye® 680RD, IRDye® 800CW) for one hour and imaged using LICOR Odyssey® Imager system.

**Murine pancreas preparation for quantification of in vivo c-Jun phosphorylation in HEI-286-KTR SCs and Trichrome staining**

5 days after injection, the pancreas was isolated and embedded in Tissue-Tek OCT (Electron Microscopy Sciences). The frozen blocks were sectioned at 10 μm thickness using a Cryostat microtome (Leica CM1950). Sections underwent DAPI staining for assessing c-Jun phosphorylation in HEI-286-KTR SCs or stained using Trichrome stain kit (Abcam, ab150686) for assessing fibrosis. Trichrome stain slides were scanned using Pannoramic 250 Flash scanner 3D (Histech). Fluorescent slides were scanned using Pannoramic Scan II scanner (3D Histech). Images were analyzed using Slide Viewer 2.6.0.166179 (3D Histech).

**Atomic force microscopy of the human PDAC sections and force curves analysis (Detailed)**

We prepared frozen-fixed samples as previously described.[3] Human PDAC samples were obtained from patients with resectable PDAC and frozen in OCT. Two days before the experiment, six 10 μm thick sections were cut with a cryostat (Leica Biosystems, CM1950). Two sections were transferred in a glass bottom Petri dish (FluoroDish FD5040), coated with poly-L-lysine (P8920 Sigma), fixed for 10 min with 4% PFA on ice, washed with cold PBS and kept on ice in 2 ml PBS until the start of the force volume AFM experiment. The measurements correspond to nerve bundles within the same section and not to same bundles across multiple sections since we have used one section per patient. The other sections were stained for H&E, P-c-Jun, GFAP, S100, and pan-cytokeratin; and used for identifications of nerves, cancer cells and stroma on the corresponding section used for AFM. Before each experiment the AFM was calibrated using the thermal noise method. The force maps were from 10x10 to 60x60 $\mu m^2$ depending on the nerve size. Force threshold for the stiffness measurement was set for 3 nN. Force curves were fitted according to the Hertz model. Data visualizations were performed using the Asylum research software.

**Atomic force microscopy on cultured cells to induce force and imaging to assess KTR distribution (Detailed)**

Cells were seeded in a glass bottom Petri dish (FluoroDish FD5040), coated with poly-L-lysine (P8920 Sigma) 48 hours prior to the experiment. Forces on cells were applied using a MFP-3D-BIO AFM microscope (Oxford Instruments) using cantilevers with 20 μm diameter borosilicate glass probes (CP-CONT-BSG-B, sQUBE) and a nominal spring constant k=0.1 N/m (Novascan). Before each experiment, the exact spring constant (varied between 300 and 400 pN/Nm) of the cantilever was determined using the thermal noise method and its optical sensitivity was determined using a PBS-filled glass bottom Petri dish as an infinitely hard surface. The cantilever was positioned over the nucleus or the cytoplasm of the cell and engaged on the cell surface with a trigger point of 120 nN. Cells were compressed for 30-40 minutes and fluorescent images were acquired every 5 minutes with a 63x 1.4 NA oil immersion objective (Carl Zeiss) using an inverted Zeiss AxioObserver Z1 microscope with the Zen Software (Carl Zeiss) and an Axiocam MRm (Carl Zeiss). For cPLA2-mKate translocation analysis, the first image was acquired 1 minute after the force was applied.

**Intravital multiphoton microscopy**

Animals were anesthetized with 5% isoflurane for 10 minutes, and once under, with 1-2% isoflurane/oxygen mixture to maintain anesthesia for the duration of imaging. Breathing patterns were continuously monitored to assess the depth of anesthesia. Animal was placed on a 35°C heating pad (Kent Scientific) and the pancreas was exposed through a 1 cm incision on the left flank. Using 10x oculars, areas positive for Clover, indicating Schwann cells, were selected for 4D imaging using a 30x silicone immersion objective in the Olympus 1200 Multiphoton-confocal hybrid system (Evident Microsystems). Multiphoton excitation at 880 nm was used, combined with filter sets for second harmonic generation (435-445 nm), and GFP (500-535 nm). 3D or 4D stacks were collected at 10-300 μm depth in z. At the end of each imaging session, the animals were euthanized, and tissues were collected. The acquired stacks were analyzed using ImageJ and Imaris software (Oxford Instruments, UK).

**Particle Image Velocimetry (PIV)**

PIV analysis was performed using PIVlab tool for MathLab[4] on collagen images of two consecutive frames of the time-lapse movie. The PIV algorithm 'Optical flow (wavelet-based)' was used. Parameters were: Vector scale 7; Vector line width 1; plot every nth vector, n=6.

**Live imaging microchannels (Detailed)**

The microchannels coated with fibronectin (10 μg/mL; Sigma-Aldrich F1141) for 1 hour at room temperature. After washing 3 times with PBS, dishes were incubated with the cell culture medium for 15 minutes at 37°C and 5% CO2 and 50,000 cells were placed in each access port. Cells in microchannel dishes were incubated at 37°C and 5% CO2 for 24 h. Migrating cells were recorded with an inverted microscope (Axio Observer, Zeiss) at 37°C with 5% $CO_2$ atmosphere and 10× and 20× objectives. Time-lapse images were taken for 24 hours every 5 or 10 minutes.

**Compression using 6-well plate confiners (Detailed)**

200,000 cells were placed on fibronectin-coated 6-well glass-bottom plates (Mattek) for 48 hours. Cells were compressed with a 6-well cell confiner (4Dcell, Cat # CSOW 620) using 2, 5, 8 or 20 μm confinement slides according to 4Dcell protocol. Cells were imaged with an inverted microscope (Axio Observer, Zeiss) and 10× or 40× objectives at 37°C with 5% $CO_2$ atmosphere.

For RNA-Seq analysis, cells were uncompressed or compressed, for 5 minutes or 4 hours, at 37°C with 5% CO2 atmosphere. The cells compressed for 5 min were recovered for 3 hours and 55 minutes at 37°C with 5% $CO_2$ atmosphere, so that RNA was extracted after 4 hours for all the samples. For immunofluorescence, cells were fixed immediately after compression using 4 % paraformaldehyde.

**RNA Sequencing Library Preparation, and Analysis**

RNA sequencing (RNA-seq) data were from three independent biological replicates. RNA was extracted from cells using the RNeasy mini kit (Qiagen, #74104) according to manufacturer's instructions. RNA-seq was performed by the MSK Integrated Genomics Operation facility. All samples underwent RiboGreen quantification Agilent BioAnalyzer quality control. 500 ng RNA of each sample were submitted for polyA selection and TruSeq library preparation using 8 PCR cycles according to Illumina instructions (TruSeq Stranded mRNA LT Kit, #RS-122-2102). After barcoding samples were run on a HiSeq 2500 in a 50 bp/50 bp paired-end run using the HiSeq SBS Kit v4 (Illumina). Sequence data processing and analysis was performed by the Bioinformatics Core at MSK. Differentially expressed genes in compressed cells were submitted for pathway enrichment analysis using EnrichR (https://maayanlab.cloud/Enrichr/).

**qPCR**

Total RNA was extracted using the RNeasy Mini Kit (Qiagen) according to the manufacturer's instructions. cDNA was synthesized from purified RNA using reverse transcription reagents (Thermo Fisher, Maxima™ H Minus cDNA Synthesis Master Mix). Quantitative real-time PCR was performed in technical triplicates using SYBR Green PCR Master Mix (Thermo Fisher, PowerTrack™ SYBR Green Master Mix) on a ViiA 7 Real-Time PCR System (Thermo Fisher). Melt-curve analysis was conducted to verify amplification specificity. Relative gene expression levels were calculated using the ΔΔCt method, with *GAPDH* serving as the endogenous normalization control. Primers used were
F 5'-GACGTGCTGGGAAGGTACAC-3' and
R 5'-AGCCCACTGTCCACTACA-3' for cPLA2 (*PLA2G4A*), and
F 5'-GAAGGTGAAGGTCGGAGTC-3' and
R 5'-GAAGATGGTGATGGGATTTC-3' for *GAPDH*.

**Inferred pathway activation/suppression (IPAS) score**

The IPAS score was computationally calculated as previously described.[5]

**Statistical analysis**

Unpaired two-sample t-tests were used for the comparison of continuous variables between two groups. Two-way ANOVA and Repeated Measures ANOVA were used for grouped analyses. Continuous variables were assessed within individual patients using a two-sided Pearson's correlation test. To analyze the intra-individual correlation between stroma stiffness and phosphorylation of c-Jun in nerves or cancer cells for cohorts of 4, 5 and 13 patients, the 'repeated measures correlation' test (rmcorr) was used [6]. Rmcorrt calculates a repeated measures correlation using a form of analysis of covariance, while controlling for "patient" as a categorical variable to account for any between-patient variation. Statistical significance was defined at P values less than 0.05. Statistical analyses were performed using Prism 8 or 10 (GraphPad Software, Inc.) and R version 4.1.1 using the 'rmcorr' package.

**Data availability**

GSE180710, GSE292211 and GSE292214 are the accession numbers for the RNA-seq data of HEI-286 cultured alone or co-cultured with MiaPaCa-2; HEI-286 under compression for 5min and 4h and not compressed; uncompressed and compressed HEI-286, MiaPaCa-2 and Panc01 for 5min, respectively.

**SUPPLEMENTARY FIGURE LEGENDS**

**Supplementary Figure 1. Stroma stiffness and SC activation markers in nerves.**

A: Correlation analysis between P-c-Jun expression (fluorescence mean intensity) in nerves and the stiffness of stroma surrounding these nerves measured by atomic force microscopy on an adjacent section for 3 individual patients (Raw data available at Source data table).

B: Correlation of P-c-Jun in nerves and corresponding stiffness of stroma surrounding these nerves using repeated measures correlation (rmcorr). The stiffness of stroma surrounding a nerve is the mean of 3 different measurements around the nerve. 11 patients. r= 0.382, P= 0.011.
C: Correlation of P-c-Jun in nerves and corresponding stiffness of these nerves using repeated measures correlation (rmcorr). The stiffness is the mean of 3 different measurements within the nerve. 13 patients. r= 0.070, P= 0.584.
D: GFAP staining in PDAC specimen showing single nerve fibers and stiffness maps of the corresponding areas indicating Young's modulus values. The indicated value of the Young's modulus is the average of 1024 values measured within one area.
E: Correlation analysis between GFAP expression (fluorescence mean intensity) in single nerve fibers and stiffness of stroma surrounding these single nerve fibers for 5 individual patients (Raw data available at Source data table).

**Supplementary Figure 2. Live monitoring of c-Jun phosphorylation in Schwann cells.**
A: The c-Jun kinase translocation reporter (KTR) contains a Jun recognition motif (JRM), phosphorylation (P) sites at nuclear localization (NLS) and nuclear export (NES) signals, and a fluorescent protein Clover domain. Phosphorylation of the JRM inhibits NLS activity and increases NES activity, inducing translocation of the fluorescent reporter from the nucleus to the cytoplasm.
B: HEI-286 SCs expressing KTR (HEI-286-KTR) allow live monitoring of c-Jun phosphorylation. When c-Jun is not phosphorylated, the fluorescent reporter is mainly localized in the nuclear (N). When c-Jun is phosphorylated, the fluorescent reporter is mainly localized in the cytoplasm (C). C/N ratio is a quantitative measure of c-Jun phosphorylation levels.
C: Anisomycin (100 ng/ml for 1 h) induce c-Jun phosphorylation in HEI-286-KTR SCs. Western-blots analysis of P-c-Jun, c-Jun and Vinculin in anisomycin and DMSO (vehicle) treated HEI-286-KTR SCs.
D: Anisomycin treatment (100 ng/ml for 1 h) induces c-Jun phosphorylation in HEI-286-KTR and HEI-286 SCs as seen by KTR translocation and P-c-Jun staining. Scale bar 20 μm.
E: c-Jun phosphorylation of three live HEI-286-KTR SCs recorded every 5 min for 20h showing periodic oscillations of the signal.

F: c-Jun phosphorylation in HEI-286-KTR SCs in presence or absence of anisomycin and JNK inhibitor SP600125 (n=20 cells per condition, repeated twice, ****$p<0.0001$, 2-way ANOVA test for comparison of anisomycin with DMSO treatments. The arrow indicates the addition of anisomycin.
G: Effect of the substrate stiffness on c-Jun phosphorylation in HEI-286-KTR SCs. (n=50 cells per condition, mean ± SD, representative of 3 experiments).
H: Effect of IL-6 on c-Jun phosphorylation in HEI-286-KTR SCs. (n=100 cells, mean ± SD, Student's t-test, repeated 3 times).
I: Effect of pancreatic cancer cells on c-Jun phosphorylation in HEI-286-KTR SCs. Human pancreatic cancer cells MiaPaCa-2 and murine pancreatic cancer cells KPC increase SC c-Jun phosphorylation (n=20 cells, mean ± SD, Student's t-test, repeated twice).
J: Heatmaps representing the difference of gene expression in HEI-286 SCs grown alone and HEI-286 cells co-cultured with pancreatic cancer cells MiaPaCa-2 (left).

**Supplementary Figure 3. Activated HEI-286-KTR cells in murine pancreas.**
A: Experimental timeline.
B: Fibrosis in pancreata from cerulein-treated mice. Masson's trichrome staining in pancreata from control and cerulein-treated mice showing increase of collagenous connective tissue fibers (blue) in cerulein-treated animals (repeated 3 times). Stiffness measurements using AFM (n=15 different locations within one section per condition, mean ± SD, Student's t-test, ** $p<0.01$, repeated 3 times)
C: Immunofluorescence staining of SOX10, EGR2 and GAP43 in fibrotic murine pancreas injected with HEI-286-KTR SCs. HEI-286 KTR SCs express the three SC markers suggesting that SCs kept a degree of SC identity.
D: Confocal microscopy of HEI-286-KTR cells in fibrotic murine pancreas 5 days after orthotopic transplantation. C/N ratio was measured for each cell and cells were classified as activated, intermediate state or non-activated cells. In activated cells, KTR-Clover is mostly localized in the cytoplasm (C/N>1.2); in non-activated cells KTR-Clover is mostly localized in the nucleus (C/N<0.8); in the intermediate state cells, KTR-Clover is distributed in both the nucleus and the cytoplasm (0.8<C/N <1.2).

**Supplementary Figure 4. Activation of HEI-286-KTR cells in the microchannels and compression chambers.**

A: Quantification of C/N ratios of NLS (red) and KTR (green) in HEI-286-KTR SCs expressing NLS and representative images. Nuclear localization of NLS before and 30 min after force, and translocation of KTR from nucleus to cytosol in the same HEI-286-KTR SCs in which 120nN forces are applied. (n=5 cells, Paired Student t test, NLS: nuclear localization signal, 120nN force applied at the cell nuclei using AFM, Paired Student t test, * indicates position of the probe at the nucleus)

B: Loss of c-Jun phosphorylation in HEI-286-KTR SCs exiting a 10, 5 and 4 µm width microchannel. Time is h:min. Scale bar: 10 µm.

C: HEI-286-KTR NLS-RFP cells in compression chambers with different pillar heights. Nucleus rupture in cells compressed in chambers with 2 µm height pillars. No nucleus rupture in compressed cells in chambers with 5 µm and 8 µm height pillars and in non-compressed cells. Scale bar 20 µm. Quantification of KTR and NLS C/N ratios. (n=5 cells per condition, mean ± SD, two ways Student t-test, *** $p<0.001$, repeated 3 times)

D: Western blot of P-c-Jun in control (Cont) and compressed (Comp, 5 µm height pillars) HEI-286 KTR cells with quantification (n=4 biological replicates, mean ± SD, two ways Student t-test, **** $p<0.0001$)

E: P-c-Jun immunofluorescence in control and compressed HEI-286-KTR SCs (compression with 5 µm height pillars, n=40 cells, mean ± SD, two ways Student t-test, **** $p<0.0001$, repeated 3 times)

F: GFAP immunofluorescence in control and compressed HEI-286-KTR SCs (compression with 5 µm height pillars, n=40 cells, mean ± SD, two ways Student t-test, **** $p<0.0001$, repeated 3 times)

**Supplementary Figure 5. Activation of HEI-286 KTR cells on different substrates.**

A: qPCR validation of the cPLA2 knockout in HEI-286 KTR SCs. (n=3, mean ± SD, two ways Student t-test, ** $p<0.01$)

B: Pathway enrichment for the upregulated genes in compressed HEI-286, MiaPaCa-2 and Panc01 cells (NCI Nature dataset).

### MOVIE LEGENDS

**Movie 1: Effect of anisomycin in HEI-286-KTR SCs.** HEI-286 SCs expressing c-Jun kinase translocation reporter Clover (HEI-286-KTR) allow live monitoring of c-Jun phosphorylation. HEI-286-KTR SCs were imaged every min. At first, the fluorescence is mainly localized in the nucleus, indicating that c-Jun is not phosphorylated. The addition of anisomycin (100ng/ml) at time 30 min induces c-Jun phosphorylation and fluorescence translocation in cytoplasm. Time presented as h:min. Scale bar 10 μm. 10 frames per second.

**Movie 2: Intravital multiphoton microscopy images of HEI-286-KTR SC squeezing through collagen-rich area.** HEI-286-KTR SC undergoes activation (fluorescence translocation from nucleus to cytoplasm) when squeezing through narrow space. Images were taken every 5 min and shown at 6 frames per second. Time is h:min:s. Scale bar 10 μm. Green: KTR-Clover, magenta: collagen.

**Movie 3: Activation and deactivation of HEI-286-KTR Clover SCs surrounded by moving collagen fibers.** Collagen fibers move toward the HEI-286-KTR SC that undergoes activation (fluorescence translocation from nucleus to cytoplasm) and not toward the HEI-286-KTR SCs that undergo deactivation. Images were taken every 5 min and shown at 2 frames per second. Time is h:min. Scale bar 10 μm. Red and blue dots mark cells undergoing activation and deactivation respectively. Green: KTR-Clover, magenta: collagen.

**Movie 4: HEI-286-KTR SCs in microchannels of different width sizes.** A HEI-286-KTR SC forms nucleus protrusions into 4 μm width microchannels and undergoes activation (fluorescence translocation from nucleus to cytoplasm). A HEI-286-KTR SC enters a 5 μm width microchannel and undergoes activation after nucleus squeezing. A HEI-286-KTR SC enters a 10 μm width microchannel without undergoing activation. Blue dots are in nuclei of not activated cells, and pink dots are in nuclei of activated cells. Time is h:min. Images are taken every 10 min. Scale bar 10 μm. 7 frames per second.
