## Supplemental Figures for "Pancreatic cancer fibrosis activates protumorigenic Schwann cells through a nuclear mechanosensing mechanism"

#### Supplementary Figure 1

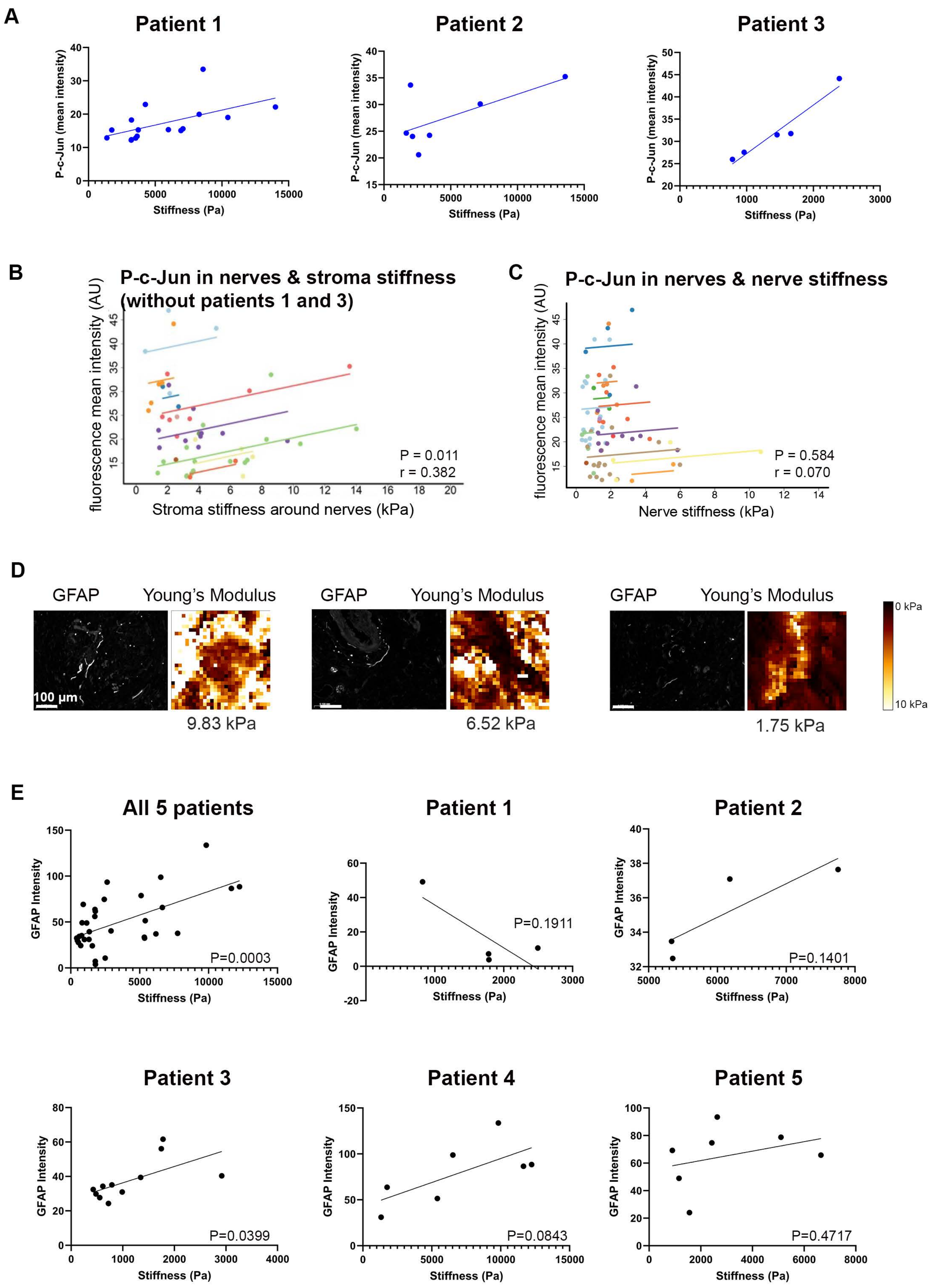

#### Supplementary Figure 2

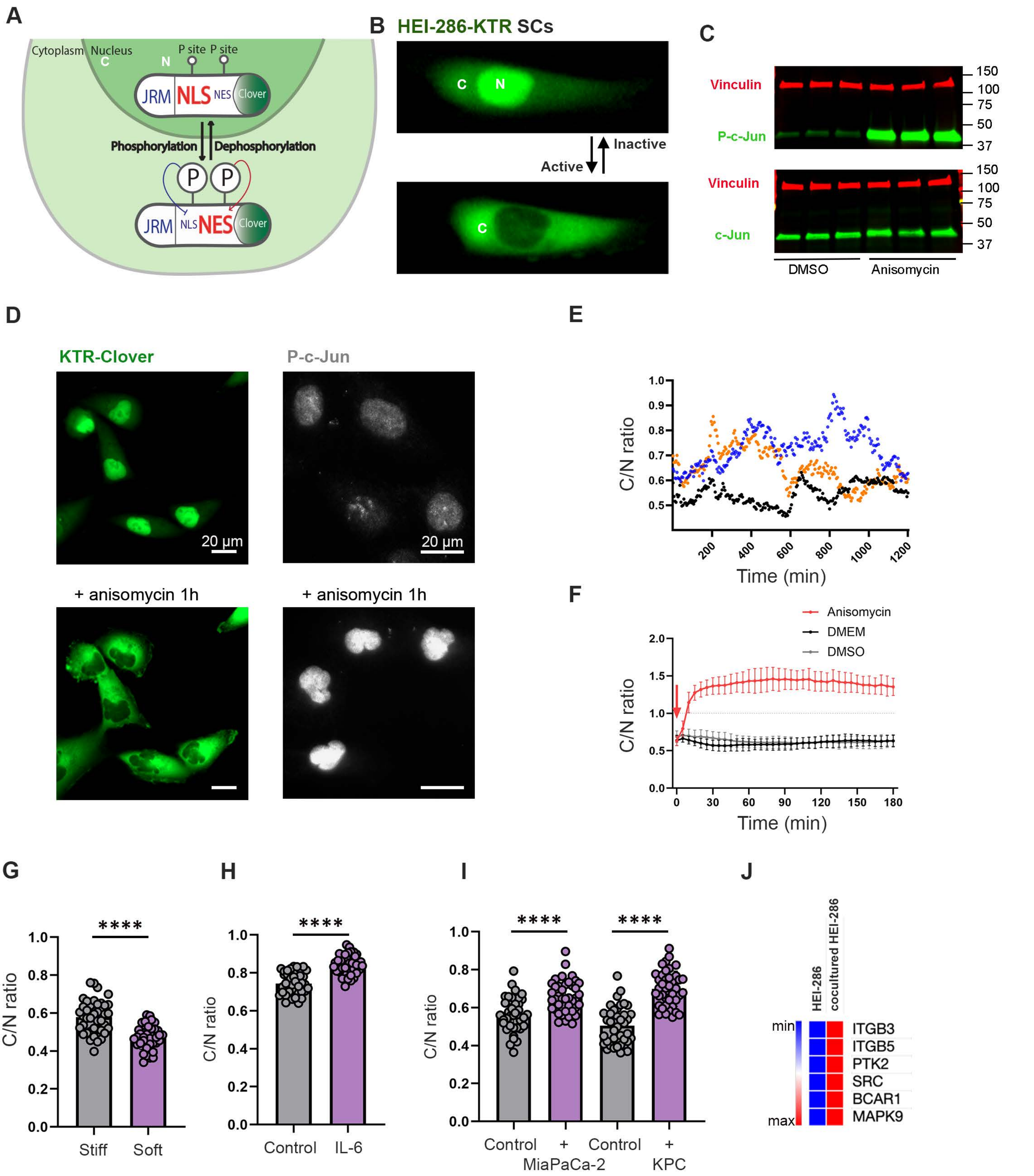

### Supplementary Figure 3

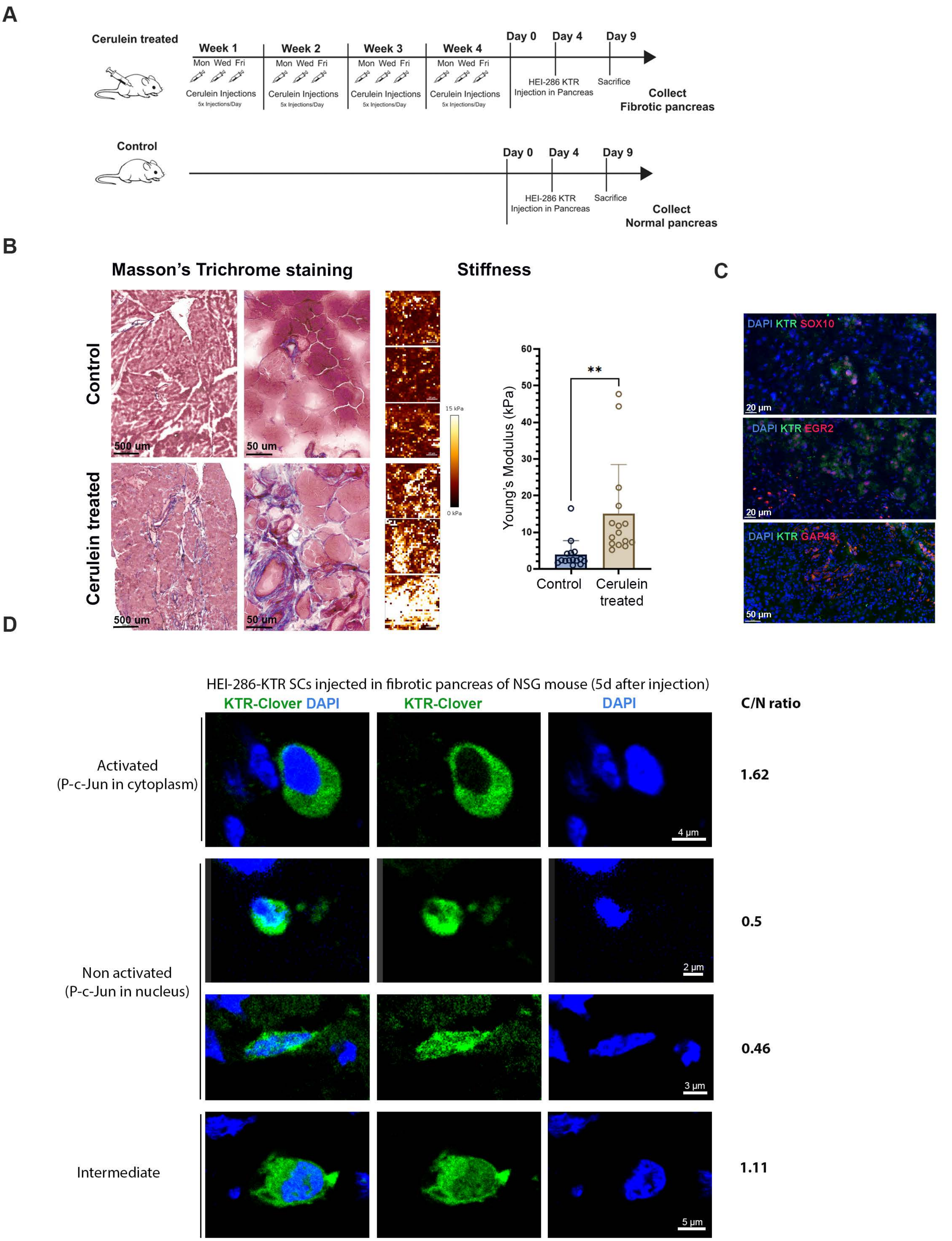

### Supplementary Figure 4

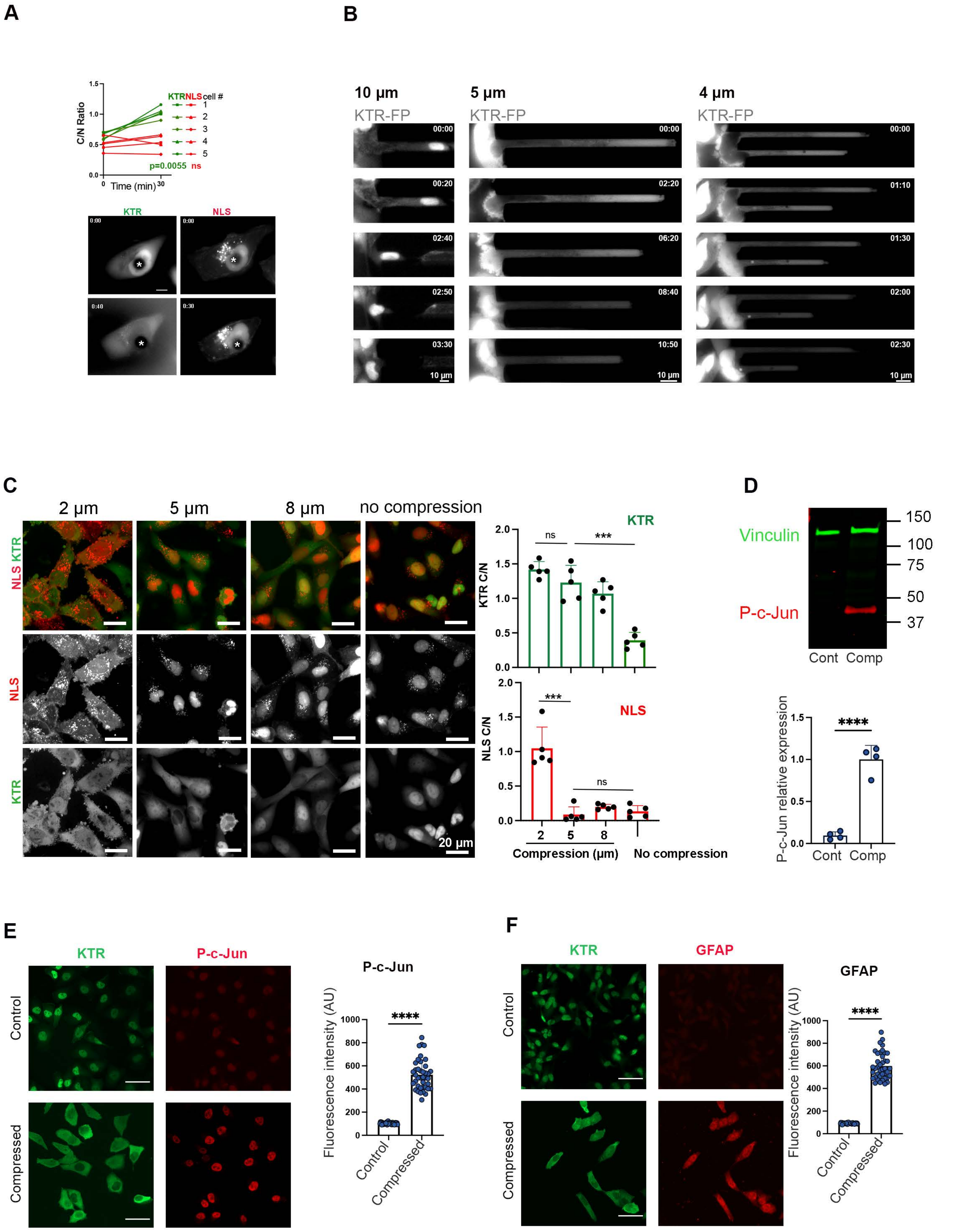

#### Supplementary Figure 5

A

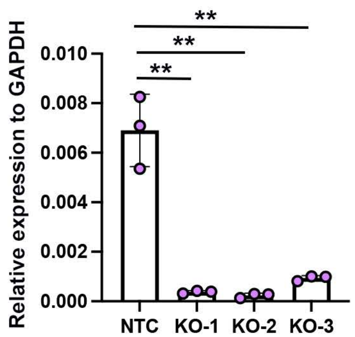


B

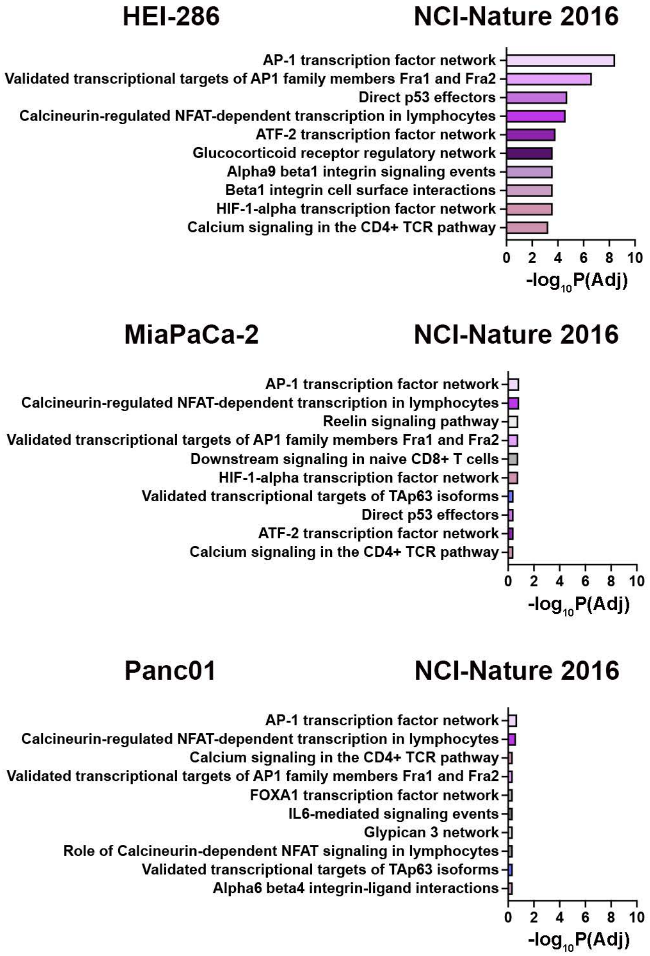
